# REST/NRSF drives homeostatic plasticity of inhibitory synapses in a target-dependent fashion

**DOI:** 10.1101/2021.04.21.440758

**Authors:** Cosimo Prestigio, Daniele Ferrante, Antonella Marte, Alessandra Romei, Gabriele Lignani, Franco Onofri, Pierluigi Valente, Fabio Benfenati, Pietro Baldelli P

## Abstract

The repressor-element 1-silencing transcription/neuron-restrictive silencer factor (REST/NRSF) controls hundreds of neuron specific genes. We showed that REST/NRSF downregulates glutamatergic transmission in response to hyperactivity, thus contributing to neuronal homeostasis. However, whether GABAergic transmission is also implicated in the homeostatic action of REST/NRSF is unknown. Here, we show that hyperactivity-induced REST/NRSF activation triggers a homeostatic enhancement of GABAergic inhibition, with increased frequency of miniature inhibitory postsynaptic currents (IPSCs) and amplitude of evoked IPSCs. Notably, this effect was only observed at inhibitory-onto-excitatory neuron synapses, whose density increased at perisomatic sites, demonstrating a strict target-selectivity. These effects were occluded by TrkB receptor inhibition and resulted from a coordinated and sequential activation of the Npas4 and BDNF gene programs. The findings highlight the central role of REST/NRSF in the complex transcriptional responses aimed at preserving physiological levels of neuronal activity in front of the ever-changing environment.

**Impact Statement:** This work elucidates the mechanisms by which the transcriptional regulator REST/NRSF selectively upregulates GABAergic transmission onto excitatory neurons in response to hyperactivity to rescue neuronal homeostasis.

## INTRODUCTION

The brain is characterized by a constant and fine homeostatic regulation of neuronal intrinsic excitability and synaptic strength aimed at keeping neuronal networks’ activity within a physiological range. The dysregulation of such homeostatic mechanisms is implicated in the early-phase progression of several neurological disorders, including epilepsy and Alzheimer’s disease (Frere and Slutsky, 2018; Lignani et al., 2020; Styr and Slutsky, 2018). An increasing amount of experimental evidence shows that the repressor element 1-silencing transcription factor (REST; also known as neuron-restrictive silencer factor, NRSF) is the molecular hub of a complex neuronal transcriptomic remodeling aimed at maintaining brain homeostasis (Zullo *et al*, 2019; Lu *et al*, 2014; Hu *et al*, 2011; Pozzi *et al*, 2013; Pecoraro-Bisogni *et al*, 2018).

REST, initially described as a nuclear negative regulator, is a zinc-finger transcription factor with several unique properties. Highly expressed in embryonic stem-cells, it is rapidly downregulated in neural progenitors and maintained at low levels after differentiation, thus enabling neurons to express the critical genes necessary for the acquisition and preservation of the neuronal phenotype (Ballas et al., 2001, 2005a; Chong et al., 1995; Ooi and Wood, 2007; Schoenherr and Anderson, 1995). REST recruits multiple corepressors, including histone deacetylases and methyltransferases, to conserved 21 bp REST binding sites, known as RE-1 sites, on target gene promoters and represses their expression in non-neuronal cells (Ballas et al., 2005b, 2001; Griffith et al., 2001). REST targets hundreds of neuron-specific genes, including those encoding for postsynaptic receptors, ion channels and transporters, cytoskeletal proteins and synaptic proteins (Bruce et al., 2004). However, some evidences showed that it may occasionally act as a transcriptional activator (Kallunki et al., 1998; Perera et al., 2015).

Although downregulated in differentiated neurons, REST levels are dynamically tuned in neurons and can be upregulated in response to kainate-induced seizures in vivo (Hu et al., 2011; McClelland et al., 2014, 2011; Palm et al., 1998; Spencer et al., 2006) and chronic hyperactivity in cultured neurons (Pecoraro-Bisogni et al., 2018; Pozzi et al., 2013). Moreover, REST levels progressively increase in cortical neurons of healthy humans over aging because of the increased Wnt/β-catenin signaling (Lu et al., 2014). While the role of REST in epileptogenesis is still debated (Baldelli and Meldolesi, 2015; Garriga-Canut et al., 2006; Hu et al., 2011; Lignani et al., 2020; McClelland et al., 2014, 2011), its function in aging was demonstrated to be protective from oxidative stress and amyloid β toxicity. Conditional deletion of REST in the mouse brain leads to age-related neurodegeneration and nuclear REST is decreased in mild cognitive impairment and Alzheimer’s brains, while elevated REST levels are associated with the preservation of cognitive functions in aged humans (Lu et al., 2014). More recently, REST was found to be upregulated in the brain of humans with extended longevity (Zullo et al., 2019).

Since neural excitation is believed to increase with age, the capability of REST to repress excitation-related genes (Pozzi et al., 2014) suggests that the activation of REST and reduction of excitatory neural activity could be responsible for slowing ageing in humans (Zullo et al., 2019). These results are consistent with our previous studies where we demonstrated that the hyperactivity-dependent activation of REST furthers neuronal-network homeostasis by down-regulating intrinsic excitability and presynaptic function of excitatory neurons in response to chronic hyperactivity (Pecoraro-Bisogni et al., 2018; Pozzi et al., 2013). However, whether REST governs homeostatic plasticity by acting also on inhibitory GABAergic interneurons is still unknown.

Here, we demonstrate that REST translocates in the nucleus of inhibitory neurons in response hyperactivity and upscales the strength and density of peri-somatic GABAergic synapses only when they contact excitatory neurons. The underlying REST-dependent mechanism for this target-specificity involves activation of the P1 promoter-regulated BDNF transcript through a REST-dependent activation of the immediate-early gene (IEG) Npas4. These findings emphasize the role of REST as a master transcriptional regulator in neurons. By specifically acting at excitatory and inhibitory connections, REST activates a coordinated program that maintains brain circuits activity within physiological levels.

## RESULTS

### Neuronal hyperactivity induces nuclear translocation of REST/NRSF in inhibitory neurons

To study the transcriptional activity of REST in excitatory and inhibitory neurons, primary hippocampal neurons from GAD67-GFP mice (Tamamaki et al., 2003; Valente et al., 2016) were treated with a Cy3-tagged decoy oligodeoxynucleotide, complementary to the RE1-binding domain of REST (Cy3-ODN), that was chemically modified to improve its stability (Soldati et al., 2011) and used to trace endogenous REST. A random decoy oligodeoxynucleotide sequence (Cy3-NEG) served as a negative control. Both Cy3-ODN and Cy3-NEG, added extracellularly, were able to efficiently cross the plasma membrane and diffuse in the cytosol (**Figure 1A,B**). Interestingly, upon hyperactivity induced by 1 h treatment with the convulsant 4AP (100 µM), Cy3-ODN doubled its partitioning ratio into the nucleus in both GFP-negative excitatory neurons and GFP-positive inhibitory neurons (**Figure 1C-E**). Conversely, Cy3-NEG treated neurons showed a homogeneous diffusion in both nucleus and cytosol that was unaffected by the 4AP treatment (**Figure 1 – figure supplement 1)**. These results confirm that REST exerts its transcriptional activity through its translocation from the cytoplasm to the nucleus also in GABAergic neurons (Shimojo, 2008; Shimojo et al., 2001; Shimojo and Hersh, 2003).

**Figure 1.**
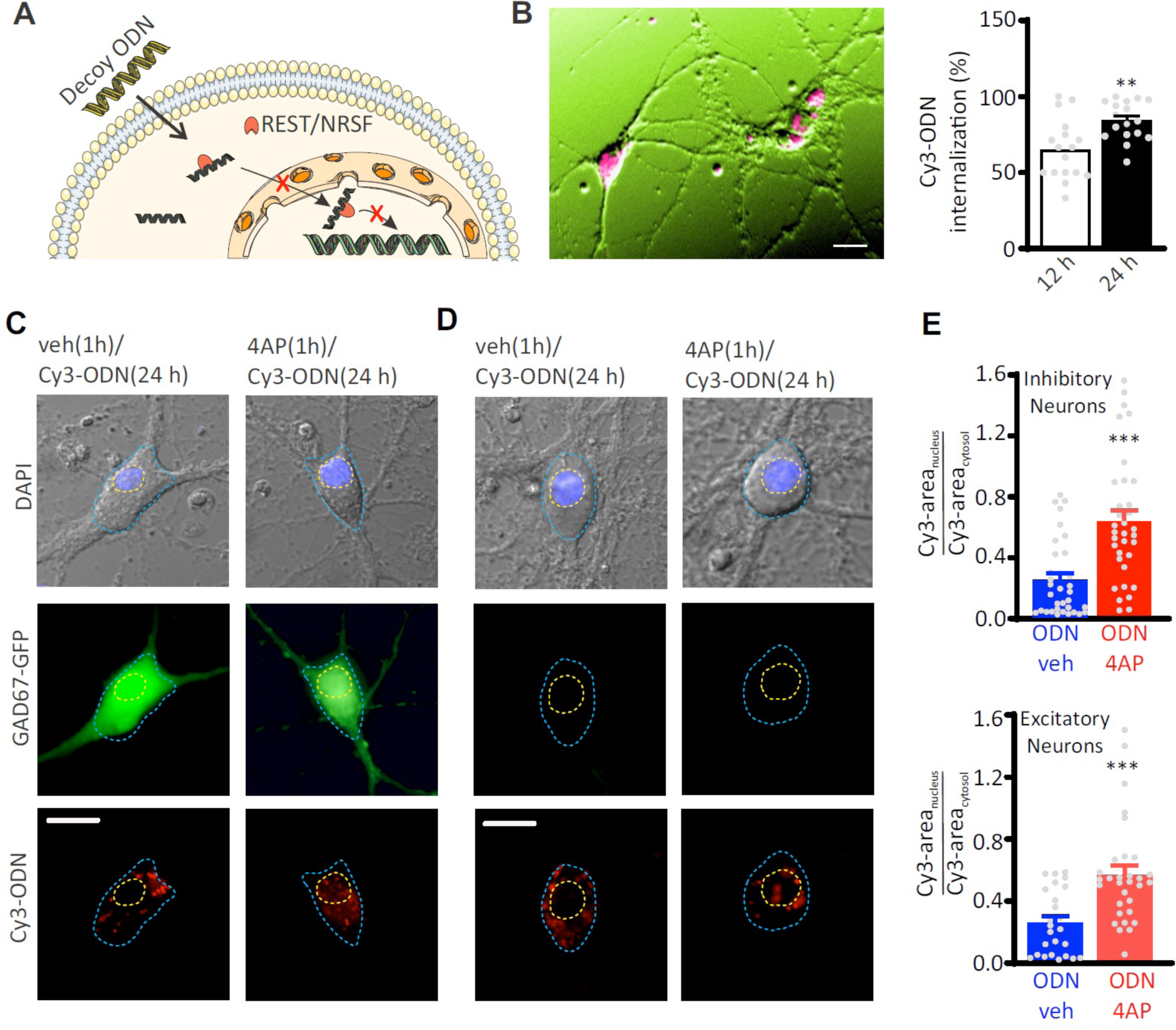
Hyperactivity induces REST translocation to the nucleus in both inhibitory and excitatory neurons. (**A**) Schematic representation of the ODN action. Having a complementary sequence to RE1, ODN binds REST, thus abrogating its capability to modulate its transcriptional activity. **(B)** *Left:* Primary hippocampal neurons (16-18 div) were incubated for 12 and 24 h with Cy3-tagged ODN (Cy3-ODN, 200 nM). *Right:* Cy3-ODN efficiently permeated into primary hippocampal neurons. The bar plot shows the mean (± sem) percentage of neurons that internalized Cy3-ODN after 12 and 24 h of incubation (n=17 fields from 2 independent preparations; **p<0.01; unpaired Student’s *t*-test). **(C-E)** Tracking of Cy3-ODN reveals both cytoplasmic and nuclear localizations of REST and its nuclear translocation upon hyperactivity in both excitatory and inhibitory neurons. **(C,D)** Representative fluorescence images of hippocampal GFP-positive inhibitory (l*eft*) and GFP-negative excitatory (*right*) neurons (17 div) treated with either vehicle or 4AP for 1 h and subsequently labeled with Cy3-ODN for 24 h. **(E)** Bar plots represent the means±sem of the REST partition ratio between cytosol and nucleus (Cy3-areanucleus/Cy3-areacytosol) in inhibitory (*upper panel*; ODN/veh n=33; ODN/4AP n=34) and excitatory (*lower panel*; ODN/veh n=25; ODN/4AP=32) neurons. ***p<0.001; Mann–Whitney’s *U*-test. Scale bars: 20 µm.

**Figure 1 - figure supplement 1.**
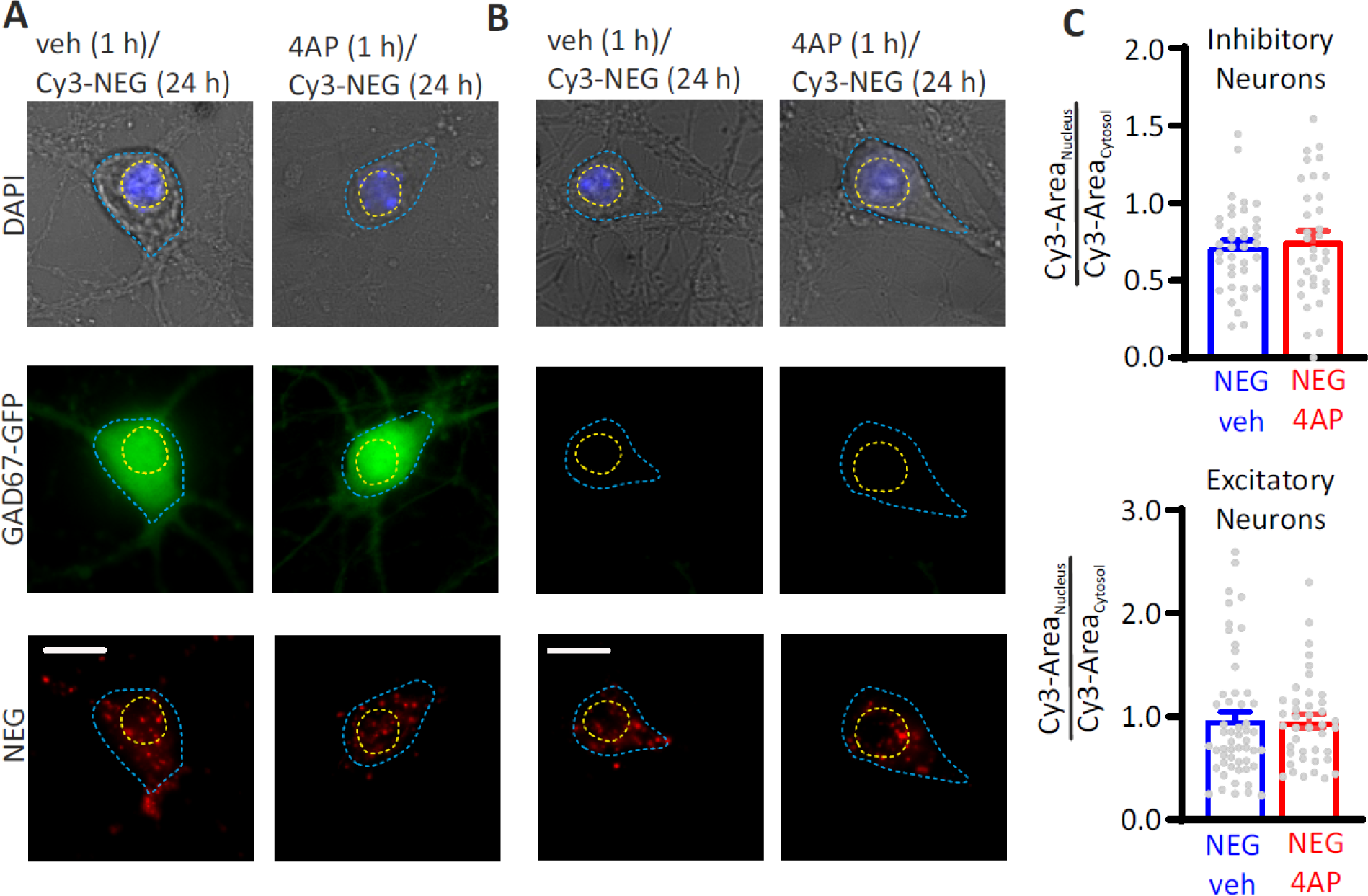
A scrambled version of ODN (Cy3-NEG) does not translocate to the nucleus on hyperactivity. Staining for Cy3-NEG revealed similar cytoplasmic and nuclear localization independently of the 4AP treatment. **(A,B)** Representative fluorescence images of hippocampal GFP-positive inhibitory (**A**) and GFP-negative excitatory (**B**) neurons treated at 17 div with either vehicle or 4AP for 1 h and subsequently labeled with Cy3-NEG for 24 h. Scale bar, 20 µm. **(C)** Bar plots (means±sem) represent Cy3-NEG partition between cytosol and nucleus (Cy3-areanucleus/Cy3-areacyto) in inhibitory (*upper panel*; NEG/veh n=38; NEG/4AP n=34) and excitatory (*lower panel*; NEG/veh n=54; NEG/4AP n=41) neurons. p=0.82 and p=0.47, respectively; Mann– Whitney’s *U*-test.

**Figure 1 – figure supplement 2.**
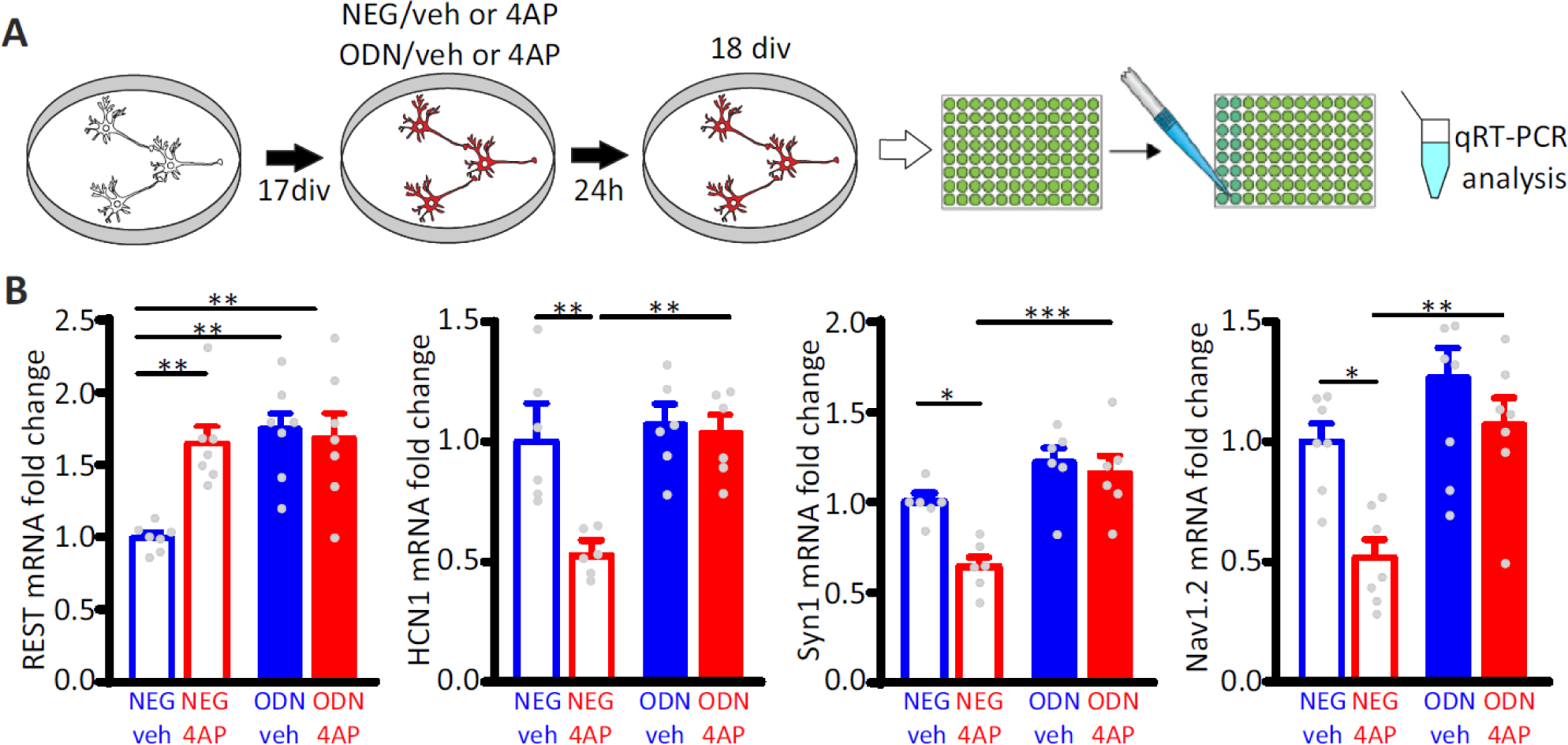
Treatment with ODN blocks the repression of REST target genes upon hyperactivity. **(A)** Schematics of the experimental protocol. Primary hippocampal neurons (16-18 div) were treated with ODN (200 nM) in the presence or absence of 4AP (100 µM) for 24 h before qRT-PCR analysis. **(B)** 4AP treatment increased REST mRNA levels, resulting in the downregulation of the REST target genes HCN1, Nav1.2 and Syn1. The effect was suppressed by ODN, demonstrating its ability to inhibit REST transcriptional silencing triggered by neuronal hyperactivity. Notably, ODN also significantly increased REST mRNA levels, suggesting the existence of REST-mediated negative control on REST transcription. Bar plots show the fold changes (means ± sem) in HCN1, Syn1 and Nav1.2 mRNAs in cortical neurons treated with NEG/vehicle, NEG/4AP, ODN/vehicle or ODN/4AP for 24 h (6<n<7 from three independent neuronal preparations). *p<0.05, **p<0,01, p<0.001; two-way ANOVA/Tukey’s tests.

The nuclear translocation of REST induced by hyperactivity was associated with downregulation of the mRNAs of HCN1, Syn1 and Nav1.2, three well-known REST target-genes (McClelland et al., 2011; Paonessa et al., 2013; Pozzi et al., 2013) in control neurons, while it was fully blocked in neurons treated with ODN (**Figure 1 – figure supplement 2**). ODN treatment also increased the transcription of REST mRNA, suggesting the presence of a feedback control of REST expression, possibly mediated by the presence of the RE1 motif within the genomic sequence of the REST gene previously reported by genome wide analysis (Johnson et al., 2007). Altogether, these data demonstrate that ODN: (i) is an excellent intracellular tracer of REST; (ii) it inhibits REST activity on its target genes; and (iii) the nuclear translocation of REST upon hyperactivity occurs in both GFP-positive inhibitory and GFP-negative excitatory neurons.

### GABAergic inhibition contributes to the REST-dependent homeostatic recovery from network hyperactivity

To investigate the role played by inhibitory transmission in REST-dependent homeostatic processes, primary cortical networks were treated at 17 days *in vitro* (div) with either NEG or ODN in the presence or absence of 4AP and their spontaneous firing activity was monitored over time (1-48 h) by microelectrode array (MEA) recordings (**Figure 2A).** In NEG-treated neurons, the mean firing and bursting rates were enhanced 1 h after the onset of 4AP stimulation with a moderate decrease of burst duration (**Figure 2B)**. Both firing and bursting rates started to decrease after 24 h of 4AP exposure and were significantly downscaled either below (firing rate) or near (bursting rate) the level of control neurons at 48 h. On the other hand, the blockade of REST activity by ODN in neurons treated for 48 h with 4AP significantly reduced the homeostatic downscaling of both mean firing and bursting rates and normalized the burst duration (**Figure 2B**).

**Figure 2.**
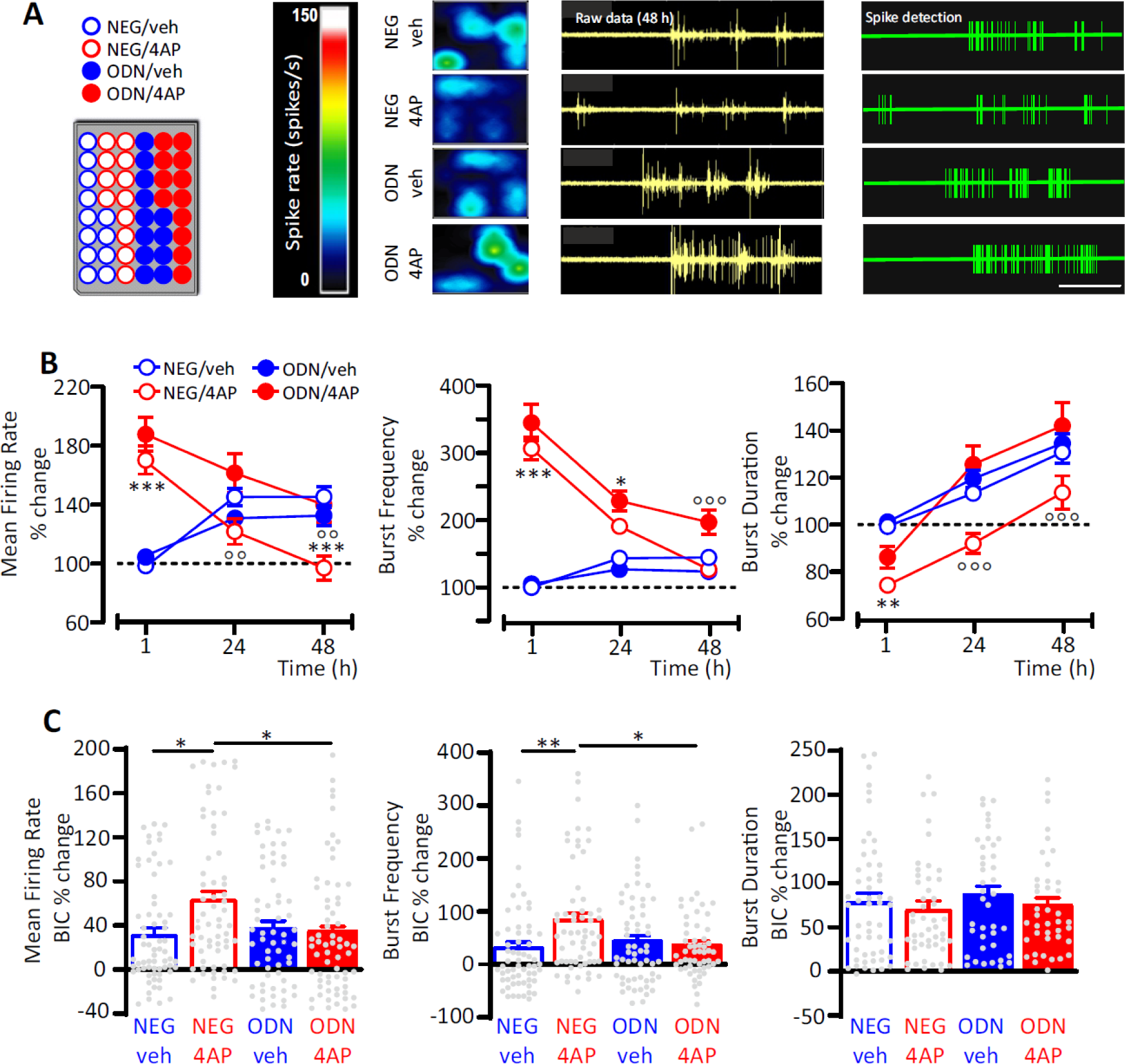
GABAergic transmission contributes to the REST-dependent homeostatic response to hyperactivity. **(A)** *Left panel:* Representative MEA plate-map showing treatment group assignment (NEG/vehicle; NEG/4AP; ODN/vehicle; ODN/4AP). *Right panels:* Firing rate heat maps (*left*), raw voltage recordings from individual electrodes (*middle*) and corresponding spike detection results (*right*), for representative wells of the four experimental groups. **(B)** Mean firing rate (*left*), bursting rate (*middle*) and burst duration (*right*) are expressed in percent of the NEG/veh baseline as a function of time after the onset of the treatments (1, 24 and 48 h). *p<0.05, ** p<0,01, ***p<0.001 NEG/veh *vs* NEG/4AP; °°p<0.01 °°°p<0.001 NEG/4AP *vs* ODN/4AP; two-way ANOVA/Tukey’s tests. (**C**) Percent increases in mean firing rate (*left*), burst frequency (*middle*) and burst duration (*right*) induced by BIC (30 μM) after 48 h of the indicated treatments. *p<0.05, **p<0.01; two-way ANOVA/Tukey’s tests. Data are means ± sem (panel B, 78<n<59; panel C, 65<n<45 MEA wells for NEG/veh, NEG/4AP; ODN/veh and ODN/4AP, from n=7 independent neuronal preparations).

To evaluate the contribution of GABAergic inhibition to the REST-dependent homeostatic response to 4AP, we blocked GABAA receptors (GABAA Rs) with bicuculline (BIC, 30 μM) at the end of the 48-h treatment with 4AP. While burst duration was similarly increased by bicuculline in all experimental groups, the GABAAR blockade induced a significantly larger increase in mean firing and bursting rates in NEG-treated neurons stimulated with 4AP than in NEG-treated neurons stimulated with vehicle (**Figure 2C**), suggesting the removal of a synaptic inhibition whose strength was increased by hyperactivity. Interestingly, such effect was abolished in ODN-treated neurons, demonstrating an involvement of REST in the upscaling of GABAergic transmission.

### Hyperactivity induces a REST-mediated increase of mIPSC frequency in excitatory, but not inhibitory, neurons

Miniature postsynaptic currents (mIPSCs) are widely used to get information about changes in synaptic properties during homeostatic plasticity (Turrigiano *et al*, 1998; Turrigiano, 2008; Pecoraro-Bisogni *et al*, 2018). While changes in amplitude mostly depend on the density or conductance of postsynaptic receptors (O’Brien et al., 1998), changes in frequency reflect the number of active synapses and/or the quantal release probability (Shao and Dudek, 2005). To evaluate whether GABAergic transmission is effectively enhanced in 4AP-treated neurons, we measured mIPSCs in excitatory (GFP-negative) and inhibitory (GFP-positive) neurons treated with NEG/vehicle, NEG/4AP, ODN/vehicle and ODN/4AP for 48 h (**Figure 3A,E**). Chronic treatment with 4AP significantly increased the frequency of mIPSCs recorded from excitatory neurons, an effect that was fully blocked by ODN (**Figure 3B,C**). Conversely, 4AP or ODN treatment did not affect the mIPSC amplitude (**Figure 3B,D**). Strikingly, 4AP-mediated chronic hyperactivity had no effects on both mIPSC frequency and amplitude recorded from inhibitory neurons (**Figure 3F-H**).

**Figure 3.**
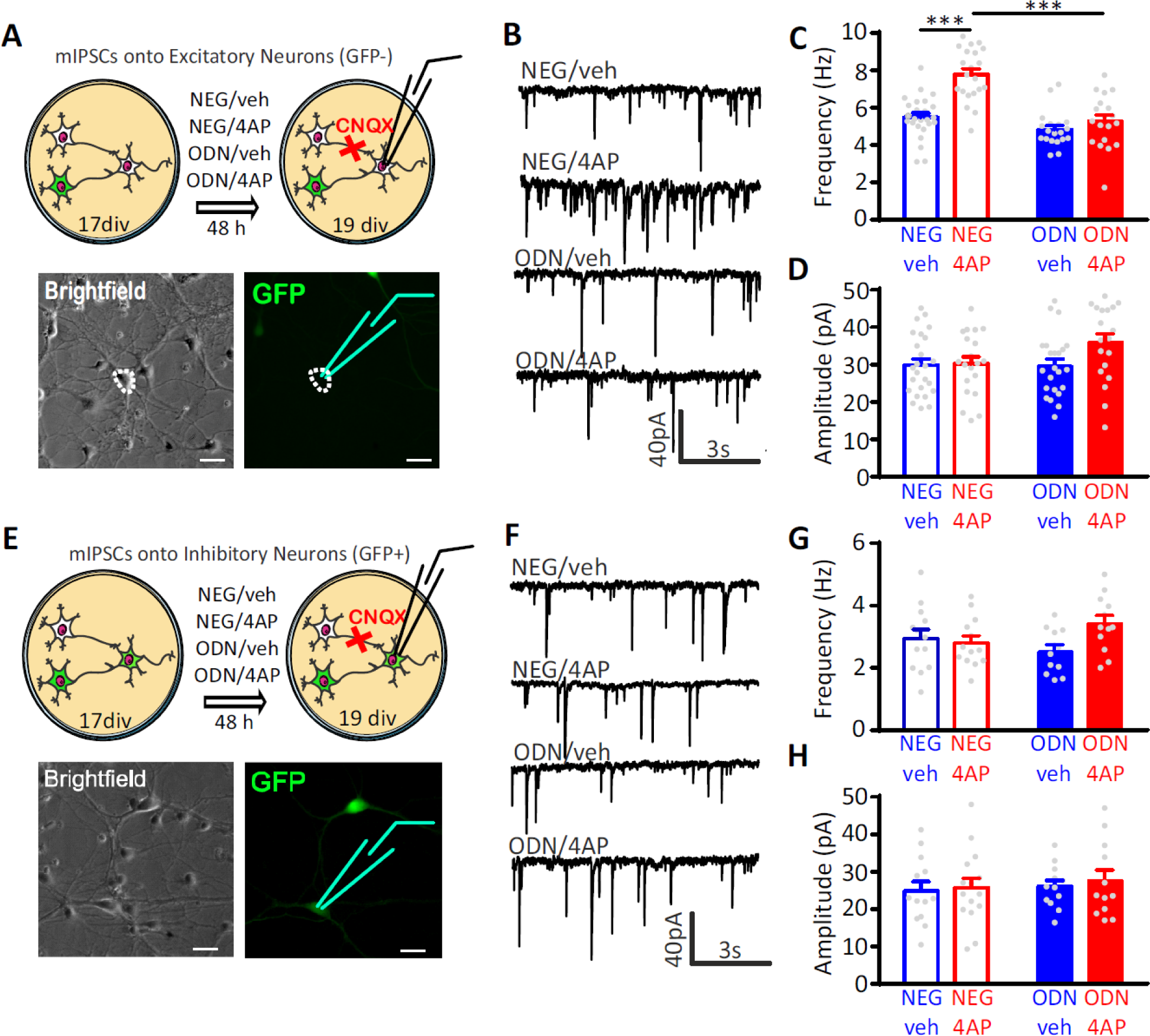
Neuronal hyperactivity selectively increases the frequency of mIPSCs in excitatory neurons in a REST-dependent fashion. (**A**) Schematic representation and representative microphotographs of a patched GFP-negative excitatory neuron. (**B-D**) Representative mIPSC traces (**B**) recorded at 19 div in the four experimental groups after 48 h of treatment. Means ± sem of IPSC frequency (**C**) and amplitude (**D**) of NEG/vehicle (n=21), NEG/4AP (n=21), ODN/vehicle (n=21) and ODN/4AP (n=21) treated neurons. (**E**) Schematic representation and representative microphotograph of a patched GFP-positive inhibitory neuron. (**F-H**) Representative mIPSC traces (**F**) recorded at 19 div in the four experimental groups after 48 h of treatment. Mean (± sem) IPSC frequency (**G**) and amplitude (**H**) of NEG/vehicle (n=13), NEG/4AP (n=13), ODN/vehicle (n=13) and ODN/4AP (n=13) treated neurons. ***p<0.001, two-way ANOVA/Tukey’s tests. Scale bars, 30 µm.

These results indicate the presynaptic origin of the REST-dependent homeostatic changes in inhibitory transmission in response to hyperactivity and demonstrate that these effects are fully target-specific, only occurring in inhibitory-to-excitatory neuron synapses.

### Neuronal hyperactivity induces a REST-mediated increase of evoked IPSCs onto excitatory neurons

To investigate whether the increased mIPSC frequency recorded in excitatory neurons is attributable to a change in the number of active inhibitory synapses or in the quantal release probability at existing sites, we investigated evoked postsynaptic inhibitory currents (eIPSCs) in 18-21 div hippocampal neurons. Extracellular minimal stimulation in loose-patch configuration was used to evoke action potentials (APs) in GFP-positive presynaptic inhibitory interneurons and patch-clamp recordings were obtained from postsynaptic GFP-negative excitatory neurons (**Figure 4A**). Evoked IPSCs in excitatory neurons treated for 48 h with 4AP displayed an increased amplitude that was suppressed upon treatment with ODN. The 4AP-induced eIPSC enhancement was accompanied by a decrease in the paired pulse ratio (PPR), that was also abolished by treatment with ODN (**Figure 4B**).

**Figure 4.**
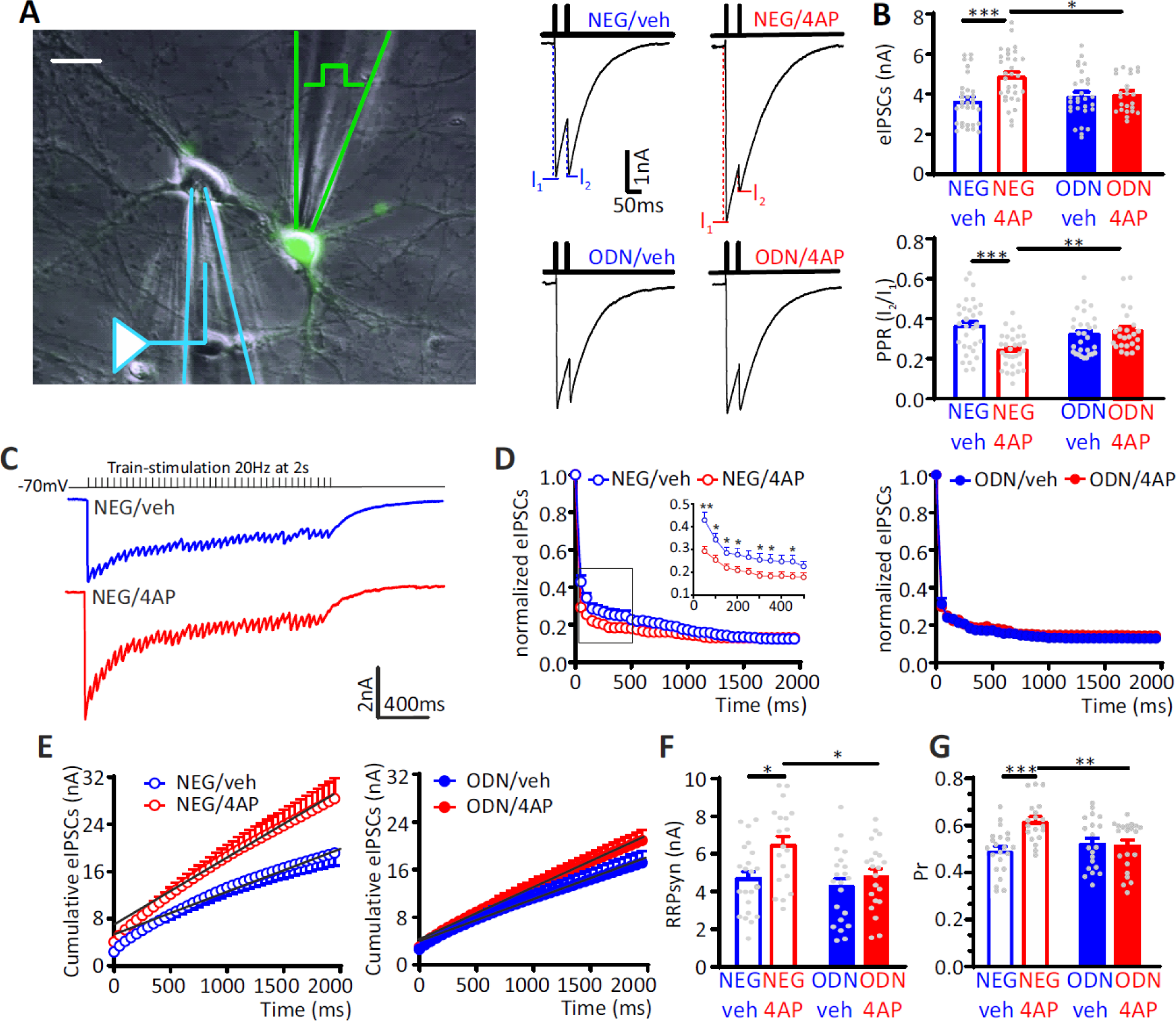
The hyperactivity-induced upscaling of eIPSCs is specific for excitatory neurons and mediated by a REST-dependent increase in RRPsyn and Pr. (A) *Left:* Experimental setup showing the stimulation electrode located on a GFP-positive GABAergic neuron and the recording electrode patching a GFP-negative excitatory neuron. Scale bar, 20 µm. *Right:* Representative eIPSCs evoked by a paired pulse stimulation protocol (interpulse interval, 50 ms). **(B)** The amplitude of the first eIPSC in the pair (*top*) and the paired pulse ratio (PPR =I2/I1; *bottom*) of NEG/vehicle (n=32), NEG/4AP (n=31), ODN/vehicle (n=29) or ODN/4AP (n=24) treated neurons are shown as means ± sem. **(C-G)** Quantal analysis of RRPsyn and Pr in GABAergic synapses onto excitatory neurons. **(C)** Representative eIPSC traces evoked by a 2 s-tetanic stimulation at 20 Hz in NEG-treated neurons in the absence (blue) or presence (red) of 4AP. **(D)** Averaged plot of normalized eIPSC amplitude *versus* time during the tetanic stimulation. The inset shows the boxed area in an expanded time scale. **(E)** Averaged cumulative profiles of eIPSCs. To calculate RRPsyn, data points in the 1-2 s range were fitted by linear regression and backextrapolated to time 0. **(F,G)** Means (±sem) of the individual values of RRPsyn (**F**) and Pr (**G**) of NEG/vehicle (n=22), NEG/4AP (n=19), ODN/vehicle (n=21) or ODN/4AP (n=21) treated neurons. *p<0.05, **p<0.01; ***p<0.001; two-way ANOVA/Tukey’s tests.

**Figure 4 – figure supplement 1.**
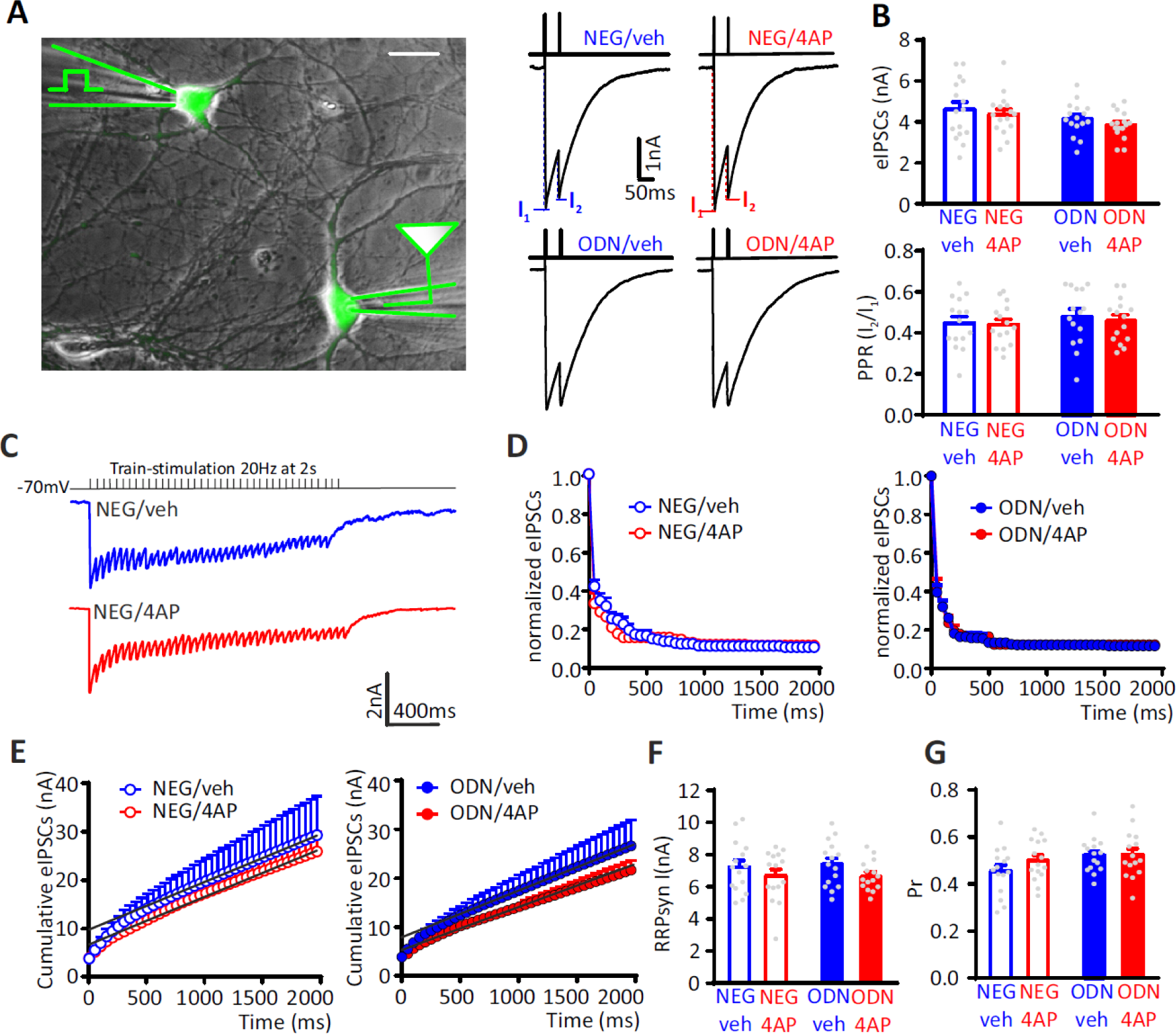
eIPSCs recorded in inhibitory interneurons are not homeostatically modulated by neuronal hyperactivity. (A) *Left:* Experimental setup showing the stimulation electrode located on a GFP-positive GABAergic neuron and the recording electrode patching a mono-synaptically connected GFP-positive inhibitory neuron. Scale bar, 20 µm. *Right:* Representative eIPSCs evoked by a paired pulse stimulation protocol (interpulse interval = 50 ms). **(B)** Lack of effect of 4AP on the amplitude of the first eIPSC in the pair *(top)* and the PPR *(bottom)* of NEG/vehicle (n=15), NEG/4AP (n=15), ODN/vehicle (n=15) and ODN/4AP (n=15) treated inhibitory neurons. Data are means±sem. **(C-G)** Quantal analysis of RRPsyn and Pr in GABAergic synapses onto inhibitory neurons. **(C)** Representative eIPSC traces evoked by a 2 s-tetanic stimulation at 20 Hz in NEG/veh- (blue) and NEG/4AP-treated neurons (red). **(D)** Averaged plot of normalized eIPSC amplitude *versus* time during the tetanic stimulation. **(E)** Averaged cumulative profiles of the eIPSCs. To calculate the RRPsyn, data points in the 1-2 s range were fitted by linear regression and back extrapolated to time 0. (**F,G**) Means (± sem) of the individual values of RRPsyn (**F**) and Pr (**G**) of NEG/vehicle (n=15), NEG/4AP (n=15), ODN/vehicle (n=15) and ODN/4AP (n=15) treated inhibitory neurons. No significant differences were found by two-way ANOVA/Tukey’s tests.

To better define the mechanism by which neuronal hyperactivity modulates GABA release, we estimated the readily releasable pool for synchronous release (RRPsyn) and the probability of release (Pr) of any given synaptic vesicle in the RRP, using the cumulative amplitude analysis (Baldelli et al., 2005; Schneggenburger et al., 1999). When inhibitory synapses onto excitatory neurons were challenged with a 2-s train at 20 Hz (40 APs), a significant depression of eIPSCs became apparent during the stimulation period, irrespective of the amplitude of the first current of the train. In NEG-treated neurons, 4AP increased synaptic depression, an effect that was virtually absent in ODN-treated cells (**Figure 4C,D)**. We next analyzed the cumulative profiles of eIPSC amplitude that display a rapid rise followed by a slower linear increase reflecting the equilibrium between depletion and constant replenishment of the RRPsyn (**Figure 4E**). The graphical extraction of the RRPsyn and Pr from the cumulative curves of individual neurons showed that the 4AP-induced eIPSC increase was due to an increase in both RRPsyn and Pr, while both quantal parameters were not affected by 4AP in ODN-treated neurons (**Figure 4F,G**).

Interestingly, the eIPSC amplitude and PPR recorded from inhibitory interneurons revealed that the synaptic strength of inhibitory synapses onto excitatory neurons was not modulated by 4AP. Accordingly, the cumulative amplitude analysis confirmed that the quantal parameters RRPsyn and Pr were unaffected by chronic neuronal hyperactivity and/or ODN treatment (**Figure 4 – figure supplement 1**). The quantal analysis of synaptic strength and short-term plasticity of inhibitory transmission in neurons exposed to chronic hyperactivity confirmed the striking postsynaptic target-specificity of the homeostatic adaptations, as already highlighted by the mIPSC analysis, demonstrating that the REST-dependent upscaling of inhibitory transmission only occurs when the postsynaptic target is an excitatory neuron.

### REST-dependent increase of peri-somatic GABAergic synapses onto excitatory neurons

Although the increased mIPSC frequency and eIPSC amplitude directly correlated with an increase in Pr and RRPsyn size, a concomitant increase in the number of functional synaptic contacts cannot be excluded. To evaluate whether the REST-mediated homeostatic response to hyperactivity also affects the density of GABAergic synapses, we triple immunostained hippocampal neurons for vGAT, gephyrin and β3-tubulin to label inhibitory presynaptic terminals, inhibitory postsynaptic scaffolds and somatodendritic structures, respectively (**Figure 5A**). We then analyzed inhibitory synapses along the soma and dendrites of GFP-negative excitatory neurons and GFP-positive inhibitory neurons. When the postsynaptic target neuron was excitatory, the sustained hyperactivity doubled the density of inhibitory perisomatic synapses and slightly, but significantly, decreased axo-dendritic contacts. The changes of both peri-somatic and axo-dendritic inhibitory synapses required an intact REST transcriptional activity, as they were fully blocked by treatment with ODN (**Figure 5B,C**). The strict target specificity of these effects was further underlined by the absence of any change on both axo-somatic and axo-dendritic inhibitory synapses when the postsynaptic target was an inhibitory neuron (**Figure 5 – figure supplement 1**).

**Figure 5.**
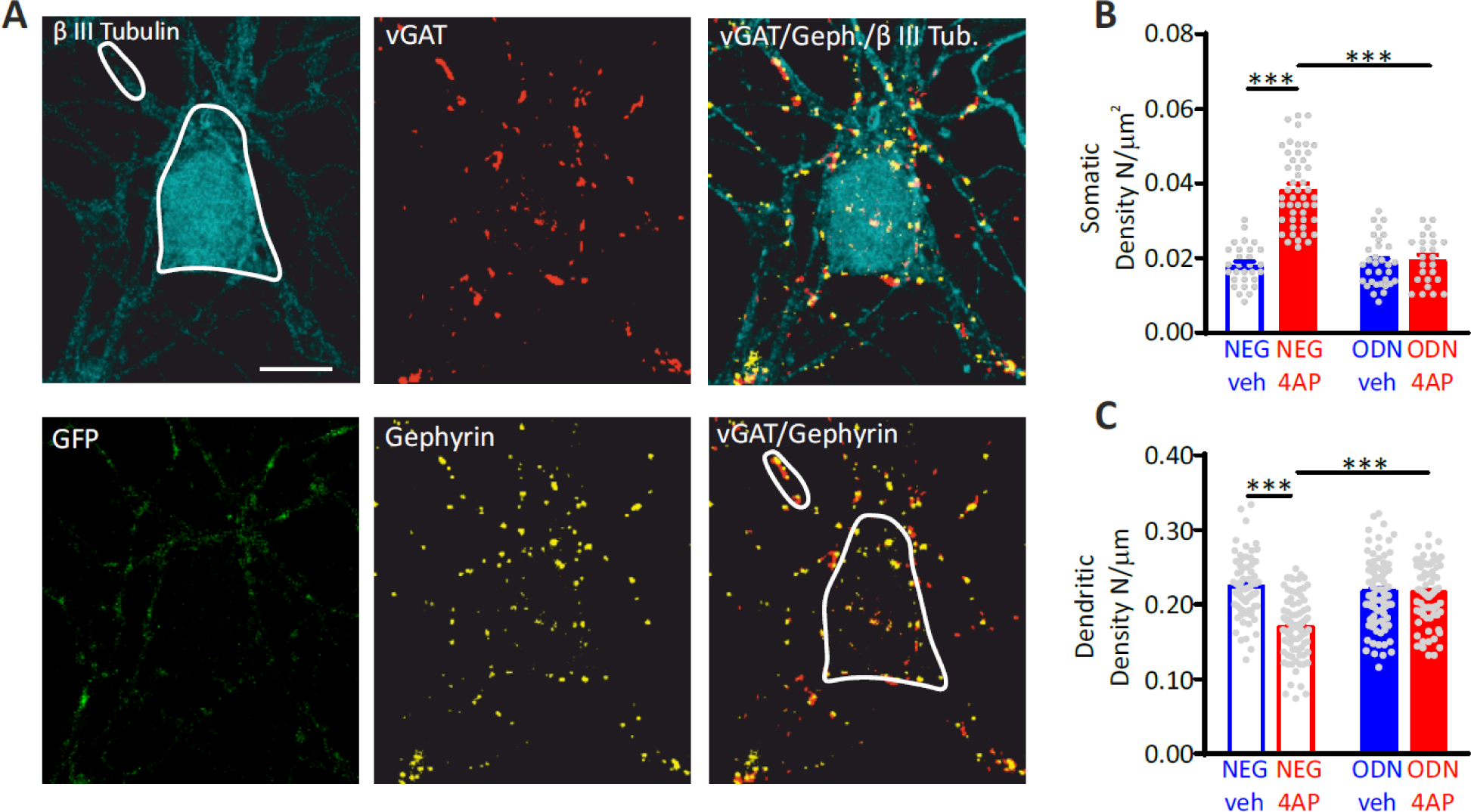
Hyperactivity induces a REST-dependent specific increase of axo-somatic inhibitory synapses onto excitatory neurons. (**A**) Representative microphotographs showing a GFP-negative excitatory neuron (20 div) labeled with β3-tubulin (light blue) and decorated with gephyrin (red) and vGAT (yellow) antibodies to identify GABAergic synapses. White lines highlight somatic and dendritic areas. Scale bar, 8 µm. (**B,C**) Quantification of the density of somatic (**B**) and dendritic (**C**) inhibitory synapses onto excitatory neurons. Data are means±sem of 48<n<25 and 87<n<71 for perisomatic and axodendritic synapses, respectively, from 3 independent neuronal preparations. ***p<0.001; two-way ANOVA/Tukey’s tests.

**Figure 5 – figure supplement 1.**
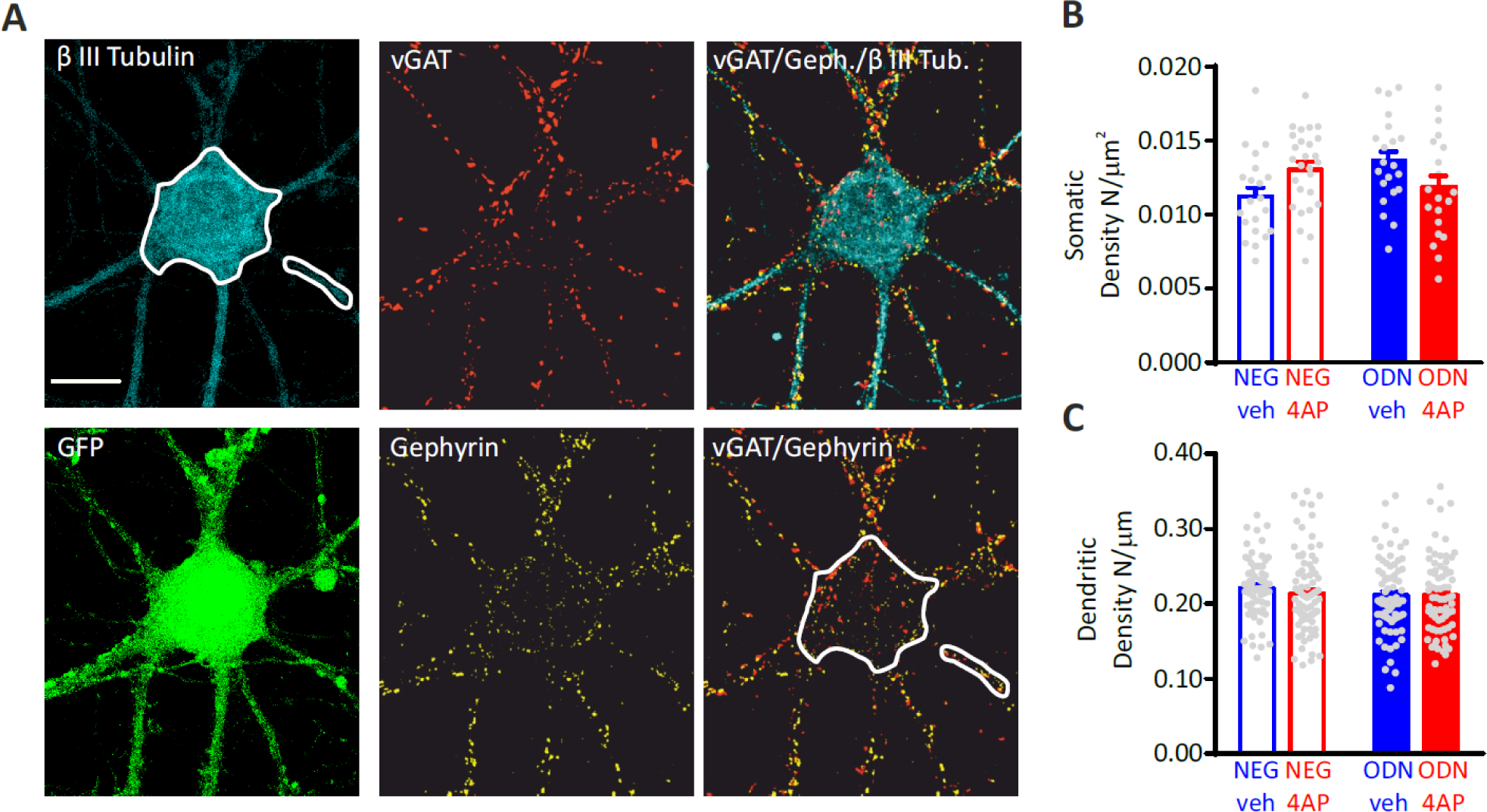
Hyperactivity does not affect the density of inhibitory synapses onto inhibitory neurons. (**A**) Representative microphotographs showing a 20 div GFP-positive inhibitory neuron labeled with β3-tubulin (light blue) and decorated by gephyrin (red) and vGAT (yellow) antibodies to identify GABAergic synapses. White lines highlight somatic and dendritic areas. Scale bar, 10 µm. (**B,C**) Quantification of the density of somatic (**B**) and dendritic (**C**) inhibitory synapses onto inhibitory neurons. Data are means±sem of 27<n<20 and 87<n<67 for peri-somatic and axodendritic synapses, respectively, from 3 independent neuronal preparations. No significant differences were found by two-way ANOVA/Tukey’s tests.

### Inhibition of BDNF-TrkB binding mimics the effect of inhibition of REST on the upscaling of inhibitory inputs onto excitatory neurons

To identify the molecular mechanisms underlying the REST-dependent and target-specific upscaling of inhibitory transmission, we investigated the possible crosstalk between REST and BDNF, suggested by the selective hyperactivity-dependent upscaling of peri-somatic inhibitory inputs to excitatory neurons. Indeed, BDNF is expressed and released primarily by excitatory neurons in an activity-dependent manner (Canals et al., 2001; Dieni et al., 2012; Ernfors et al., 1990; Hofer et al., 1990; Matsumoto et al., 2008), has a well-known action on the functional development of GABAergic synapses (Huang *et al*, 1999; Marty *et al*, 2000; Seil & Drake-Baumann, 2000; Yamada *et al*, 2002; Ohba *et al*, 2005; Lin *et al*, 2008), mainly acting at the presynaptic level (Baldelli et al., 2005, 2002). In addition, experimental evidence demonstrated that BDNF acts on the inhibitory inputs by selectively strengthening perisomatic GABAergic synapses (Fiorentino *et al*, 2009; Jiao *et al*, 2011). We first analyzed the time-course of the changes in REST and coding sequence-BDNF (cds-BDNF), mRNA levels in neurons treated with 4AP for 6, 24, 48 and 96 h (**Figure 6A**). We also included the mRNA of the membrane trafficking protein Synaptotagmin-4 (Syt4), known to localize in BDNF-containing vesicles and modulate their release (Dean et al., 2009). A significant increase in REST, cds-BDNF and Syt4 mRNA levels was already apparent after 6 h of treatment with 4AP. While the REST elevation persisted for 48 h and recovered to basal level only after 96 h, the fast increments in cds-BDNF and Syt4 mRNAs were short-lasting and fully recovered their basal values after 48 h (**Figure 6A**).

**Figure 6.**
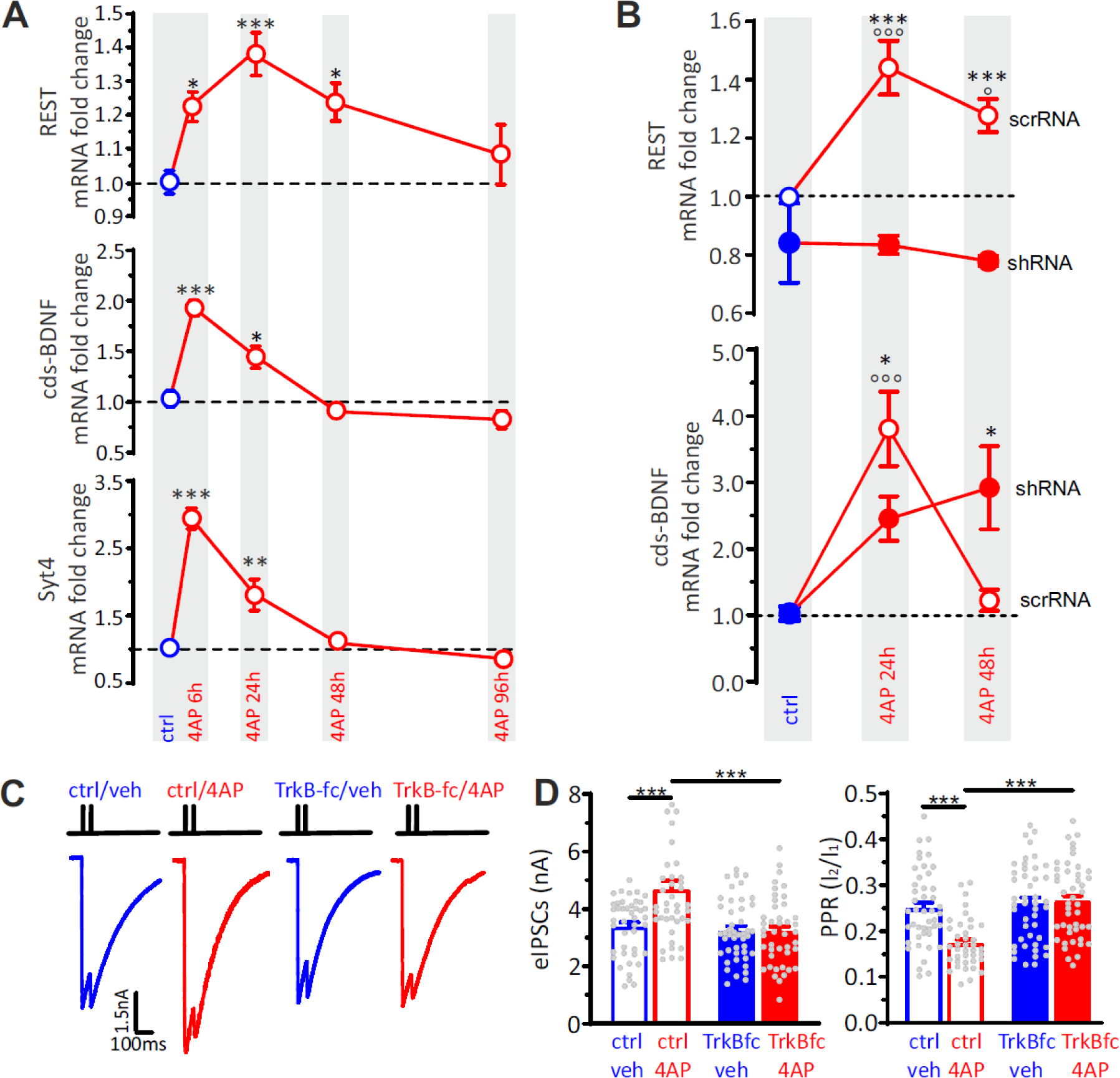
BDNF specifically potentiates GABAergic synapses onto excitatory neurons in response to hyperactivity. (**A**) Time-course of REST, cds-BDNF and Syt4 mRNA fold changes in control (blue symbols) and 4AP-treated (red symbols) cortical neurons treated for various times with 4AP. All 4AP-treated samples were collected at 20 div after 6, 24,48 and 96 h of 4AP treatment, respectively. The control sample was also collected at 20 div without any prior treatment. For each time point, data are means±sem of 9<n<10 from 3 independent neuronal preparations. *p<0.05, **p<0.01, ***p<0.001 *vs* control; one-way ANOVA/Dunnett’s tests. (**B**) RT-qPCR analysis of changes (means±sem) of REST and cds-BDNF mRNA transcript levels in control (blue symbols) and 4AP-treated (red symbols; 24 and 48 h) neurons that had been infected with either scrRNA (*open symbols*) or shRNA (*closed symbols*) viruses. Treatments were performed as described in A, with neurons collected at 20 div. For each time point, data are means±sem of 9<n<10 from 3 independent neuronal preparations. *p<0.05, ***p<0.001 shRNA *vs* scrRNA; °p<0.05, ^ooo^p<0.001 4AP-treated *vs* control; two-way ANOVA/Tukey’s tests. (**C**) Representative eIPSCs onto GFP-negative excitatory neurons in response to paired-pulse stimulation recorded in ctrl/veh, ctrl/4AP, TrkB-fc/veh and TrkB-fc/4AP treated neurons. (**D**) Mean (±sem) amplitude of the first eIPSC (I1; left) and paired-pulse ratio (PPR =I2/I1; right) of ctrl/veh (n=43), ctrl/4AP (n=39), TrkB-fc/veh (n=44) and TrkB-fc/4AP (n=43) treated neurons. *p<0.05, ***p<0.001, two-way ANOVA/Tukey’s tests.

To investigate the molecular relationship between REST and BDNF transcription, we compared the hyperactivity-dependent increase of REST and BDNF mRNAs in control and REST knockdown (KD) hippocampal neurons (**Figure 6B**). While the hyperactivity-dependent increase of REST mRNA was fully suppressed by REST KD (Pecoraro-Bisogni et al., 2018; Pozzi et al., 2013), the temporal profile of cds-BDNF mRNA induced by hyperactivity was markedly altered by REST silencing. REST deficiency significantly reduced the transient increase of BDNF at 24 h, while it completely suppressed the recovery of BDNF mRNA to basal levels at 48 h, transforming the bell-shaped BDNF expression curve into a monotonic increment.

To evaluate whether BDNF could contribute to the REST-dependent upscaling of inhibitory synapses, the effect of 4AP on evoked inhibitory transmission was studied in hippocampal excitatory neurons (18 div) treated for 48 h with either vehicle (ctrl) or the BDNF-scavenger TrkB/fc (TrkB/fc), a recombinant chimeric protein that suppresses BDNF binding to TrkB receptors (Sakuragi et al., 2013) (**Figure 6C**). Interestingly, the previously observed increase of eIPSC amplitude and decrease of PPR evoked by 4AP were fully suppressed by BDNF sequestration (**Figure 6D**).

In line with this result, the 4AP-induced enhancement of perisomatic GABAergic contacts onto excitatory neurons was also fully suppressed in neurons treated with TrkB/fc (**Figure 7A,B**). On the contrary, the 4AP-induced decrease of dendritic GABAergic contacts to excitatory neurons was insensitive to BDNF sequestration (**Figure 7A,C**). Finally, as previously shown in neurons treated with ODN, either 4AP-induced hyperactivity or BDNF scavenging did not affect the density of both axo-somatic and axo-dendritic inhibitory boutons when the postsynaptic target was an inhibitory neuron (**Figure 7 – figure supplement 1**).

**Figure 7.**
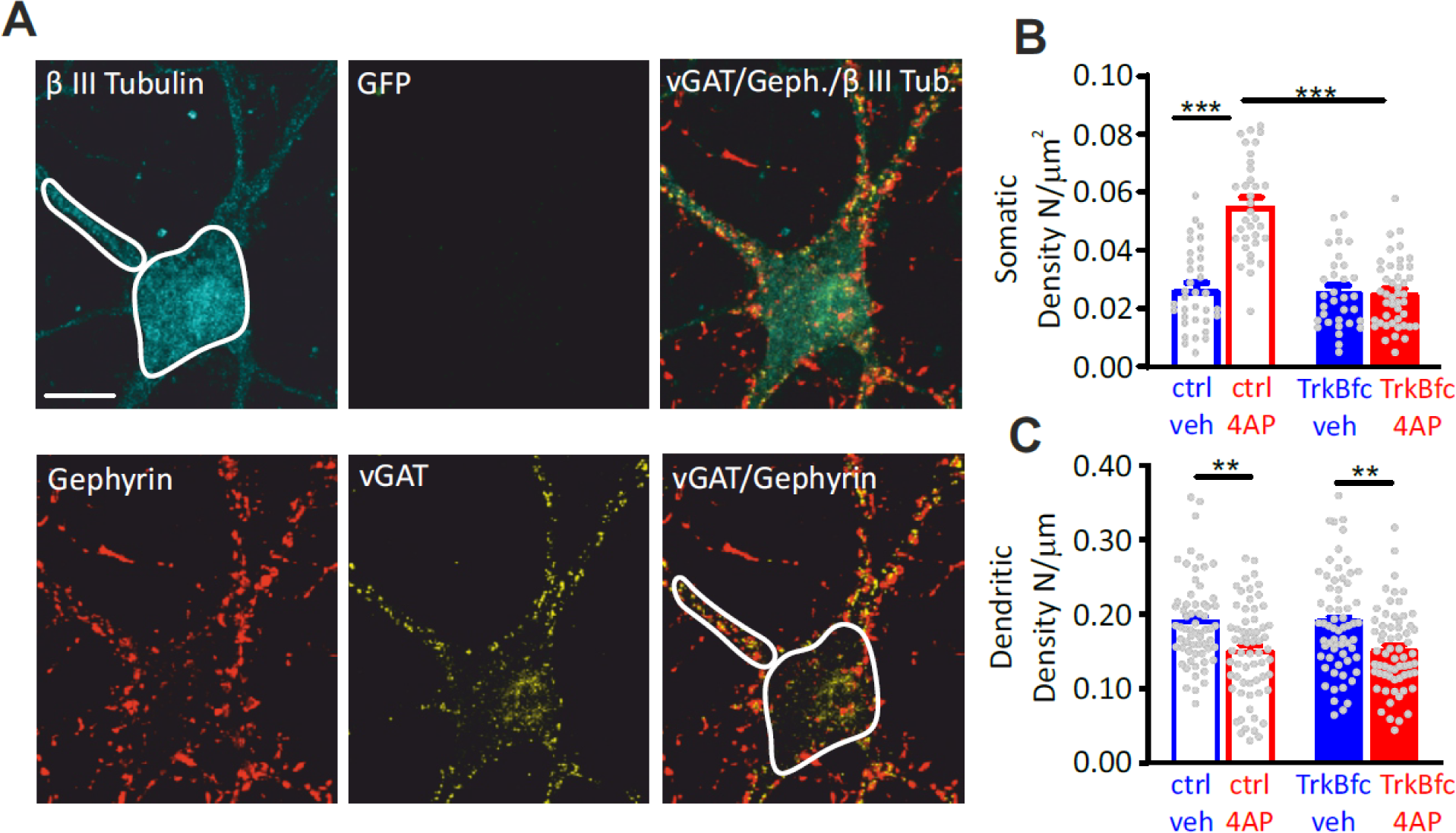
BDNF scavenging suppresses the 4AP-induced enhancement of perisomatic GABAergic contacts to excitatory neurons. (**A**) Representative microphotographs showing a GFP-negative excitatory neuron (20 div) labeled with β3-tubulin (light blue) and decorated by gephyrin (*red*) and vGAT (*yellow*) antibodies to visualize GABAergic synapses. White lines highlight somatic and dendritic areas. Scale bar,15 µm. (**B,C**) Quantification (means±sem) of the densities of somatic (**B**) and dendritic (**C**) inhibitory synapses onto excitatory neurons (33<n<44 and 63<n<64, respectively, from 2 independent neuronal preparations). ***p<0.001; two-way ANOVA/Tukey’s tests.

**Figure 7 – figure supplement 1.**
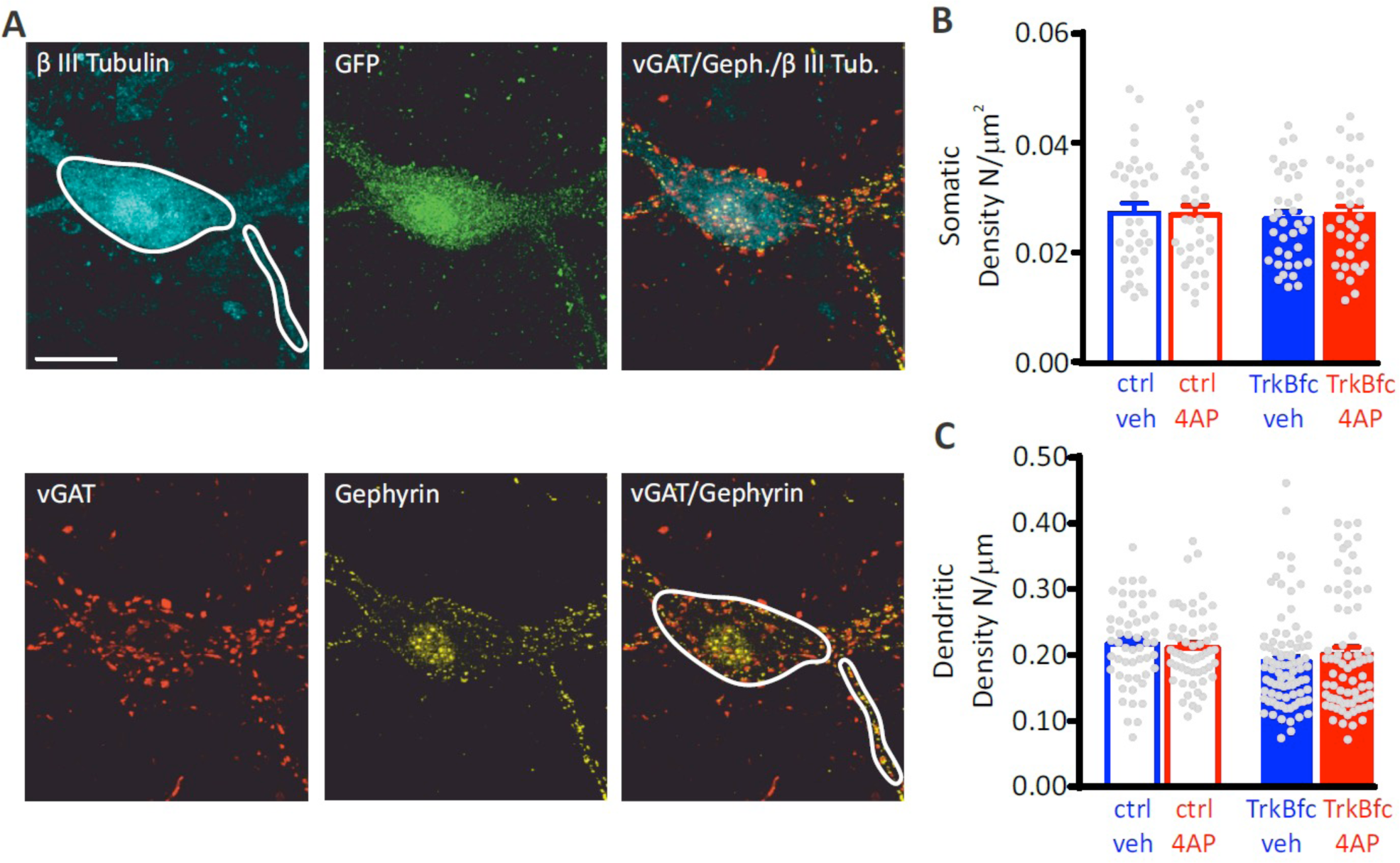
The lack of effect of 4AP on the density of inhibitory synapses onto inhibitory neurons is not affected by BDNF. (**A**) Representative microphotographs showing a 20 div GFP-positive inhibitory neuron labeled with β3-tubulin (light blue) and decorated by gephyrin (*red*) and vGAT (*yellow*) antibodies to identify GABAergic synapses. White lines highlight somatic and dendritic areas. Scale bar, 15 µm. Quantification of the density of somatic (**B**) and dendritic (**C**) inhibitory synapses onto inhibitory neurons in ctrl/veh, ctrl/4AP, TrkB-fc/veh and TrkB-fc/4AP treated neurons. Data are means±sem of 33<n<36 and 53<n<85 for peri-somatic and axodendritic synapses, respectively, from 2 independent neuronal preparations. No significant differences were found by two-way ANOVA/Tukey’s tests.

### REST triggers the activation of the “Npas4-BDNF gene program”

We have shown that REST exerts a temporal constraint on the increase of BDNF mRNA, allowing its full recovery to basal levels after 48 h (see Figure 6B). This effect was expected and widely supported by experimental evidence showing that BDNF is a REST-target gene (Garriga-Canut *et al*, 2006; Paonessa *et al*, 2016; Hara *et al*, 2009; Otto *et al*, 2007; Bruce *et al*, 2004).

On the contrary, the increase of BDNF mRNA observed in neurons treated for 24 h with 4AP was significantly reduced in REST KD neurons, demonstrating that REST plays an unexpected role in the early increase of BDNF induced by hyperactivity (see Figure 6B). Searching for a possible causal link between REST and BDNF, we focused our attention on Npas4, a transcriptional activator recently demonstrated to induce a hyperactivity-induced increase in the number and strength of perisomatic inhibitory synapses onto excitatory neurons by inducing BDNF release from excitatory neurons (Bloodgood et al., 2013; Lin et al., 2008; Spiegel et al., 2014).

The possible crosstalk between REST, Npas4, cds-BDNF and Syt4 was investigated by analyzing the changes in their transcripts in cultured neurons treated with NEG/vehicle, NEG/4AP, ODN/vehicle and ODN/4AP at a higher temporal resolution (0, 1, 3, 6, 12, 24 h). Interestingly, the 4AP-induced increase of REST mRNA was very fast, peaking at 1 h and persisting for 24 h (**Figure 8A**). As previously mentioned, ODN treatment *per se,* rapidly increased REST mRNA levels under basal conditions (see Figure 1 – figure supplement 2; Johnson *et al*, 2007), demonstrating that the basal levels of REST protein exert a tonic transcriptional repression on the REST gene.

**Figure 8.**
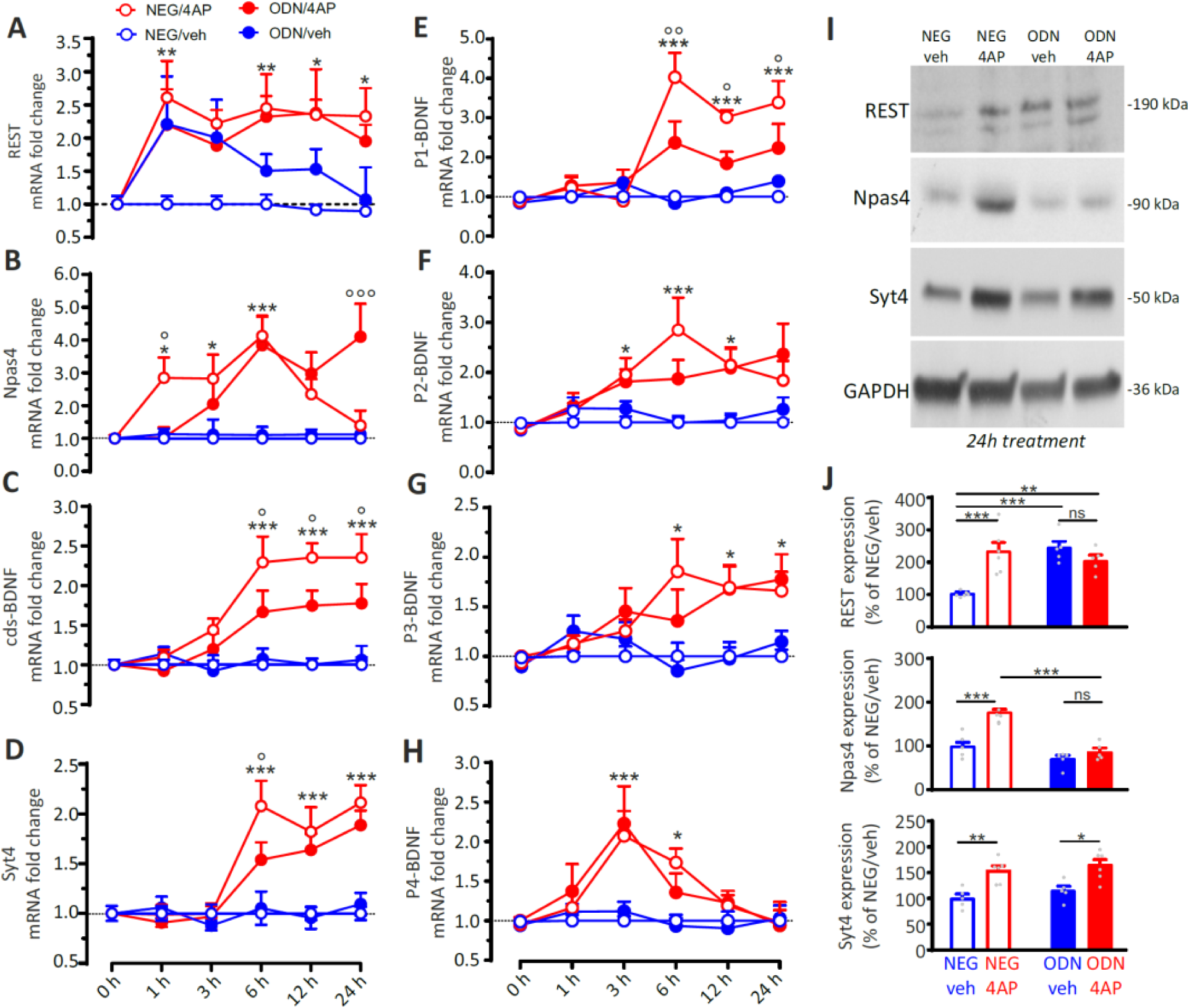
The REST-dependent potentiation of GABAergic synapses involves Npas4 and P1-BDNF activation. (**A-H**) Time course of the REST-dependent transcriptomic profile induced by hyperactivity. Mean (± sem) fold changes of REST (**A**), Npas4 (**B**), cds-BDNF (**C**), Syt4 (**D**), P1-BDNF (**E**), P2-BDNF (**F**), P3-BDNF (**G**) and P4-BDNF (**H**) mRNAs in cortical neurons treated with NEG/vehicle, NEG/4AP, ODN/vehicle or ODN/4AP before and at various times after the respective treatments. All values are normalized to the NEG/vehicle mRNA levels. For each time point, 11<n<5 from 3 independent neuronal preparations. *p<0.05, **p<0.01, ***p<0.001, NEG/veh *vs* NEG/4AP; °p<0.05, ^oo^p<0.01, ^ooo^p<0.001, NEG/4AP *vs* ODN/4AP; two-way ANOVA/Tukey’s tests. (**I,J**) Representative immunoblots (I) and quantitative analysis (L) of REST, Npas and Syt4 protein expression in cortical neurons treated for 24 h with NEG/vehicle, NEG/4AP, ODN/vehicle or ODN/4AP. All values (means±sem) are normalized to the NEG/vehicle level. GAPDH was included as control of equal loading. For each protein, 6<n<5 from 3 independent neuronal preparations. *p<0.05, **p<0.01; two-way ANOVA/Tukey’s test.

As previously reported (Lin et al., 2008), Npas4 mRNA raised already 1 h after 4AP, but the increase was transient, and its levels started to decrease at 12 h to return to basal values after 24 h. Inhibition of REST activity by ODN delayed the Npas4 rise and suppressed the return of Npas4 to basal levels (**Figure 8B**). These results demonstrate that REST affects Npas4 transcription kinetics in a manner similar to that shown for BDNF transcription (see Figure 6B), but developing on a shorter time scale. These data suggest that, while REST plays a role in the early induction of Npas4 mRNA, the sustained increase in REST levels could favor the return of Npas4 mRNA to basal levels. Moreover, 4AP-induced hyperactivity also promoted increases in cds-BDNF and Syt4 mRNAs that were delayed (6h) with respect to the raise of REST and Npas4 transcripts (1h); this delayed rise in cds-BDNF and Syt4 mRNAs was also attenuated by REST inhibition with ODN treatment (**Figure 8C,D**) Since BDNF in one of the main actors in the development of GABAergic inputs, the fine spatial and temporal regulation of the expression of distinct BDNF-transcripts could contribute to the REST-dependent potentiation of peri-somatic GABAergic inhibition. The mRNAs changes of P1-, P2-, P3, and P4-BDNF splice variants, known for their high expression in the postnatal hippocampus and cortex and for their activity-dependent regulation (Aid et al., 2007) were studied in cultured neurons treated with NEG/vehicle, NEG/4AP, ODN/vehicle and ODN/4AP for different times. While all BDNF transcripts were increased by hyperactivity (**Figure 8E-H**), only the P1-BDNF mRNA was significantly inhibited by ODN (**Figure 8E**). This specific effect is highly suggestive, in view of the previously demonstrated crucial role that Npas4 selectively exerts on the activity-dependent transcription of P1-BDNF (Lin et al., 2008; Pruunsild et al., 2011).

To confirm the REST-dependent transcriptional changes induced by neuronal-hyperactivity at the protein level, cultured neurons (17 div) treated with NEG/vehicle, NEG/4AP, ODN/vehicle and ODN/4AP for 24 h were processed by western blotting (**Figure 8I**). This analysis, confirmed the previously observed increase of REST expression in response to 4AP treatment (Pecoraro-Bisogni et al., 2018; Pozzi et al., 2013). On the other hand, the similar increase of REST protein induced by ODN, independently of the presence of 4AP, further confirms the feedback regulation by REST on its own expression (see Figure 1 – figure supplement 2B; Figure 8A). In parallel, 24 h of hyperactivity induced an increase of both Npas4 and Syt4 protein levels. However, when 4AP was applied in the presence of ODN, the increase of Npas4 protein was fully suppressed, while Syt4 overexpression was unaffected (**Figure 8J**).

Taken together, the data suggest a sequential mechanisms in which the target-specific enhancement of inhibitory transmission induced by hyperactivity involves the activation of the BDNF transcript controlled by the P1 promoter (Aid et al., 2007) through a REST-dependent activation of Npas4, a fast immediate early gene known for its ability to induce the homeostatic potentiation of somatic inhibitory inputs onto excitatory target neurons in response to neural hyperactivity (Bloodgood *et al*, 2013; Spiegel *et al*, 2014).

### The REST-dependent effects are associated with increased expression of specific markers of GABAergic transmission

We finally analyzed the changes in the transcripts encoding for GAD65, vGAT, GAD67 and the ε-subunit of GABAARs (**Figure 9A-D**) over the same timescale adopted for the investigation of the transcriptional profiles of REST, Npas4 and BDNF (0,1,3,6,12,24 h). While GAD65, the enzyme that mediates the transient GABA synthesis (Kash et al., 1997), was not modulated by hyperactivity,

**Figure 9.**
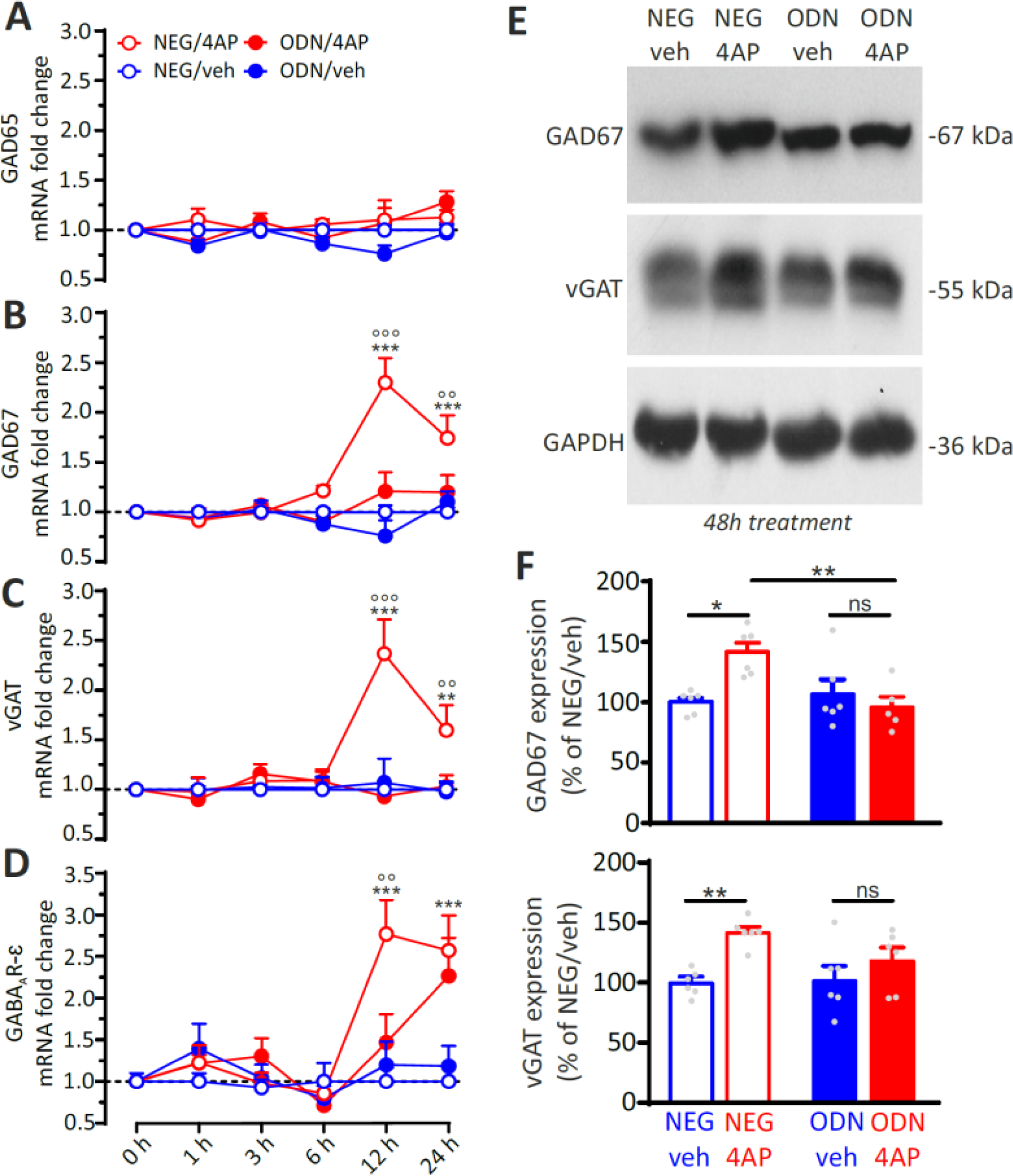
REST-dependent activation of GABAergic synaptic genes. (**A-D**) Time course of the fold changes in mRNA levels of GAD65 (**A**), GAD67 (**B**), vGAT (**C**) and GABAAR Ɛ-subunit (**D**) in cortical neurons treated with NEG/vehicle, NEG/4AP, ODN/vehicle or ODN/4AP before and various times after the respective treatments. All values (means±sem) are normalized to the NEG/veh levels. For each time point, 9<n<6 from 3 independent neuronal preparations. **p<0.01, ***p<0.001 NEG/veh *vs* NEG/4AP; °°p<0.01, °°°p<0.001 NEG/4AP *vs* ODN/4AP; two-way ANOVA/ Tukey’s tests. (**E,F**) Representative immunoblots (**E**) and quantitative analysis (**F**) of GAD67 and vGAT protein expression in cortical neurons treated for 48 h with NEG/vehicle, NEG/4AP, ODN/vehicle or ODN/4AP. All values (means±sem) are normalized to the NEG/vehicle level. GAPDH was included as control of equal loading. For each protein, 6<n<5 from 3 independent neuronal preparations. *p<0.05, **p<0.01; two-way ANOVA/Tukey’s tests.

GAD67, which regulates the constitutive synthesis of 90% of neuronal GABA, as well as vGAT and GABAAR ε-subunit showed significant increases of their mRNAs under 4AP stimulation (**Figure 9A-D**). Interestingly, while the REST, Npas4 and BDNF mRNA elevations peaked within 6 h from the onset of hyperactivity, (see Figure 8A-H), the mRNA increase of these terminal target genes was temporally shifted and peaked at 12 h. However, while the increase in transcription of the “presynaptic” genes vGAT and GAD67 was fully blocked by ODN, the postsynaptic GABAAR ε-subunit mRNA showed a slower increase that was not clearly sensitive to REST activity blockade (**Figure 9A-D**). These results were confirmed by immunoblot analysis (**Figure 9E**) that revealed a significant increase of both GAD67 and vGAT protein levels in response to hyperactivity that was suppressed when 4AP was applied in neurons treated with ODN (**Figure 9F**).

## DISCUSSION

We previously showed that neuronal hyperactivity causes REST upregulation that is crucial for maintaining neural network homeostasis through downscaling of intrinsic excitability and excitatory synaptic transmission (Pozzi *et al*, 2013; Pecoraro-Bisogni *et al*, 2018). The recently reported REST upregulation in the brain of subjects with extended longevity (Zullo et al., 2019), further emphasizes the central role of the excitatory-inhibitory balance of neural circuits, not only in the control of brain excitability, but also in the regulation of lifespan. In this paper, we investigated the homeostatic role of REST at the level of GABAergic synapses challenged by long-term treatment with the convulsive agent 4AP, the same experimental paradigm used in the studies on excitatory transmission (Pecoraro-Bisogni *et al*, 2018).

To uncover the role of REST in the hyperactivity-induced changes of GABAergic transmission, we used sequestration of endogenously expressed REST by ODN. In contrast with shRNAs that inhibit REST activity by slowly reducing its expression (Pecoraro-Bisogni et al., 2018; Pozzi et al., 2013), ODN operates a fast silencing of both constitutively and newly expressed REST, permitting a better temporal dissection of the effects of REST activation by hyperactivity (McClelland et al., 2014, 2011). REST translocated to the nucleus in response to sustained hyperactivity in both excitatory and inhibitory neurons. However, in inhibitory neurons it promoted transcriptional effects that were opposite to those elicited in excitatory neurons and strictly specific for the postsynaptic target. This resulted in a potentiation of the strength of inhibitory synaptic connections at the presynaptic level, accompanied by a structural increase in the density of perisomatic inhibitory synapses. Both effects were exclusively occurring when the postsynaptic target was an excitatory, but not an inhibitory, neuron. Network electrophysiology with MEAs testified that the REST-dependent potentiated GABAergic inhibition is an important component of the homeostatic recovery from hyperactivity and helps networks decrease the excitatory tone and re-establish a physiological excitation-inhibition balance.

The observed REST-dependent presynaptic changes affected the transcriptional profile of inhibitory neurons with increased transcription of the presynaptic markers GAD67 and vGAT and strengthened the functional and structural connectivity between presynaptic inhibitory neurons and postsynaptic excitatory targets with increased (i) frequency of mIPSCs, (ii) density of perisomatic inhibitory synapses, (iii) amplitude of eIPSCs and (iv) RRP size and release probability. Although these transcriptional and functional changes affected presynaptic inhibitory neurons, their actuation depended on the interaction with the target excitatory neuron. This implies that the REST-dependent transcriptional cascades activated by hyperactivity involve a retrograde crosstalk between excitatory neurons and inhibitory presynaptic terminals, restricting the synaptic changes only to the inhibitoryàexcitatory synaptic contacts.

This retrograde crosstalk prompted us to investigate the possible relationships between REST and BDNF. BDNF is known for its capability to strengthen GABAergic inputs (Huang et al., 1999; Marty et al., 2000; Seil and Drake-Baumann, 2000), particularly at axo-somatic synapses to excitatory neurons (Jiao et al., 2011) by acting at the presynaptic level (Baldelli et al., 2005, 2002; Valente et al., 2012). Moreover, BDNF is expressed and released in response to hyperactivity mainly by excitatory neurons, and only at low or undetectable levels by interneurons (Hofer et al., 1990; Matsumoto et al., 2008). Thus, BDNF could be the downstream effector of the target-specific homeostatic effects of REST on inhibitory transmission. Indeed, a link between REST and BDNF was experimentally demonstrated by the following observations: (i) the inhibition of BDNF-TrkB binding fully recapitulated the effects of REST inhibition on strength and density of perisomatic inhibitory synapses onto excitatory neurons; (ii) the knockdown of REST reduced the increase in the cds-BDNF transcript induced by hyperactivity; (iii) the progressive temporal shift between the prompt (1 h) elevation of REST expression, the activation of cds-BDNF transcription (6 h) and the final effects on transcription of the presynaptic GABAergic markers GAD67 and VGAT (12 h).

Interestingly, REST had opposite time-dependent effects on the BDNF response to hyperactivity, with an early enhancement followed by a late inhibition that confined the BDNF response within a limited time window. While the latter action is coherent with the transcriptional repressive action of REST on BDNF expression (Bruce et al., 2004; Garriga-Canut et al., 2006; Hara et al., 2009; Otto et al., 2007; Paonessa et al., 2016), the early enhancement of the BDNF response by REST is somewhat unexpected and may involve an indirect effect. In this respect, it has been shown that the occupancy of RE-1 sites on the promoters of REST target genes can occasionally induce transcriptional activation (Kallunki et al., 1998; Perera et al., 2015), favoring the expression of other transcription factors. Relevant to this study, it has been reported that the deletion of the RE1 element in the Npas4 promoter exerted a negative effect on Npas4 transcription (Bersten et al., 2014).

Npas4 is one of the most recently identified IEGs with peculiar characteristics (Lin et al., 2008). It is among the most rapidly induced IEGs, it is neuron-specific and selectively activated by neuronal activity (Sun and Lin, 2017). Moreover, Npas4 is known to upregulate peri-somatic inhibitory synapses to excitatory neurons, an effect mediated by the activity-dependent transcription of BDNF (Bloodgood *et al*, 2013; Spiegel *et al*, 2014; Lin *et al*, 2008) that mimics the homeostatic effects of REST described here. This makes Npas4 the most likely candidate in mediating the effects of REST on the upscaling of inhibitory inputs in response to hyperactivity. When considering the complex temporal links between REST, Npas4, BDNF and Syt4 responses after the onset of hyperactivity, both REST and Npas4 mRNAs increased after 1 h of heightened activity, while BDNF and Syt4 mRNAs displayed a later increase. The data indicate that REST, similar to Npas4,(Lin et al., 2020; Sun and Lin, 2017) can also be considered a fast IEG. Moreover, it is possible that REST, located upstream of Npas4, exerts the double effect of permitting a very early Npas4 response and of limiting the Npas4 activation window in time. The REST-Npas4 interplay has a sequential impact on cds-BDNF and Syt4 mRNAs, whose peaks at 6 h were REST-dependent. It is known that the various BDNF promoters are differentially responsive to neuronal activity (Aid et al., 2007). While the P4 transcript responded more quickly, the P1 transcript, which showed the highest upregulation, was the only REST-dependent one. This result further supports the existence of a sequential REST-Npas4 mechanism, since Npas4 is known to specifically control the activity-dependent activation of P1-BDNF transcription (Pruunsild et al., 2011).

Taken together, the data suggest that REST is an IEG that exerts a permissive role in the activation “Npas4/P1-BDNF” gene transcription cascade and has a crucial role in the temporal confinement of the transcriptional response to hyperactivity. It is tempting to speculate that the REST-dependent homeostatic upscaling of inhibitory inputs to counteract hyperactivity develops through three sequential transcriptional waves. The initial wave is characterized by the activation of fast IEGs such as REST and Npas4, in which REST has an early facilitating role on Npas4. A second transcriptional wave involves the expression of the P1-BDNF transcript by Npas4, potentially restricted to the soma of excitatory neurons (Brigadski et al., 2005; Dean et al., 2012; Pattabiraman et al., 2005). In turn BDNF, secreted by the excitatory neurons, retrogradely reaches the inhibitory presynaptic terminals where it activates, through the TrkB signaling pathway, the final transcriptional wave of effector genes that actuates the homeostatic synaptic changes (**Figure 10**). In conclusion, the data strengthen the idea that REST is a master transcriptional regulator able to interact with other transcription factors in a coordinated manner and promote both intrinsic and synaptic homeostasis in response to the variety of stressors that continuously perturb the physiological level of activity of brain circuits.

**Figure 10.**
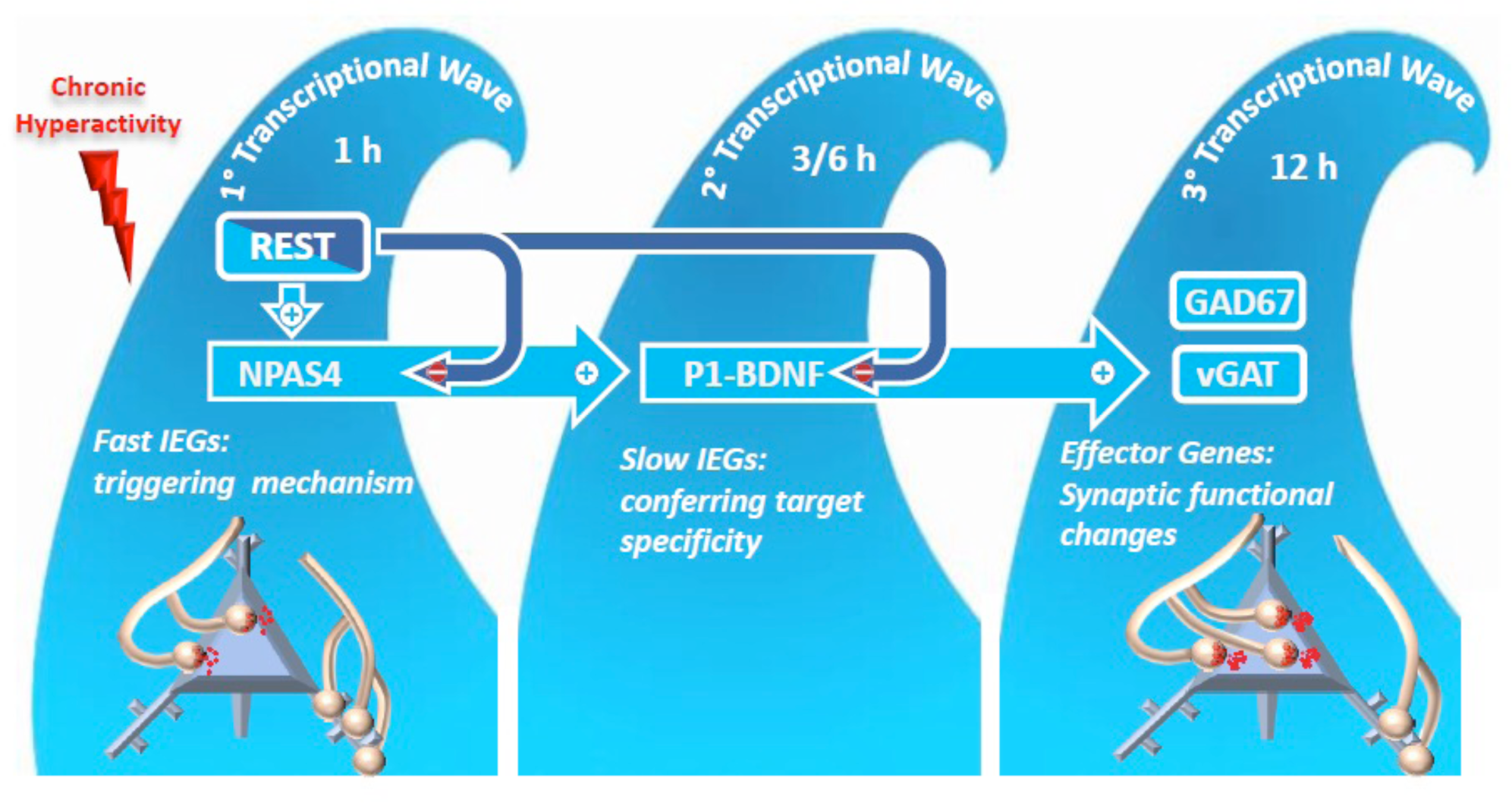
Mechanistic model of REST-dependent homeostatic upscaling of inhibitory inputs in response to hyperactivity. The process develops through three sequential transcriptional waves. The “*1^st^ transcriptional wave*” characterized by the initial activation of the IEGs REST and Npas4, whereby REST has a precocious facilitating role on Npas4. The following “*2^nd^ transcriptional wave*” that involves the expression of the P1-BDNF transcript by Npas4 that potentially restricts the functional changes to the soma of excitatory neurons. Finally, BDNF secreted by the excitatory neurons retrogradely reaches the inhibitory terminals. In these neurons, BDNF activates, via the TrkB signaling pathway, the final *“3^rd^ transcriptional wave”* of effectors genes (vGAT and GAD67), that increases the strength of axo-somatic inhibitory inputs to excitatory target neurons through a presynaptic mechanism.

## MATERIALS AND METHODS

### Key resource table

3 Key resource table
| Reagent type (species) or resource | Designation | Source or reference | Identifiers | Additional Information |
| --- | --- | --- | --- | --- |
| Experimental model | GAD67-GFP | Tamamaki et al., 2003 | n/a | n/a |
| Experimental model | C57BL6/J | Charles River | n/a | n/a |
| Chemical compound | Trypsin | Gibco | Cat. #1505014 | 0,25% |
| Chemical compound | HBSS | Gibco | Cat. # 14170-088 | n/a |
| Chemical compound | HEPES | AppliChem | Cat. #A3724,0500 | 10 mM |
| Chemical compound | D-Glucose | Sigma-Aldrich | Cat. #G7021 | 30 mM |
| Chemical compound | Gentamicin solution | Sigma-Aldrich | Cat. #G1272 | 5µg/ml |
| Chemical compound | KOH | AppliChem | Cat. # 131515 | Water solution 1M |
| Chemical compound | Bovine serum Albumin (BSA) | Sigma-Aldrich | Cat. #A4503 | 10% |
| Chemical compound | MgSO<sub>4</sub> + H<sub>2</sub>O | Sigma-Aldrich | Cat. #63138 | 6 mM |
| Chemical compound | Poly-L-lysine hydrobromide | Sigma-Aldrich | Cat. #P2636 | 0,1 mg/ml |
| Chemical compound | Boric Acid | Sigma-Aldrich | Cat. #B-7901 | For Borate Buffer 0,1M |
| Chemical compound | di-Sodium tetraborate decahydrate | Sigma-Aldrich | Cat. #A267508 | For Borate Buffer 0,1M |
| Chemical compound | Neurobasal-A Medium | Gibco | Cat. #10888-022 | n/a |
| Chemical compound | Neurobasal Medium | Gibco | Cat. # 21103049 | n/a |
| Chemical compound | B-27 Supplement | Gibco | Cat. #17504044 | 2% |
| Chemical compound | GlutaMAX™ -I (100X) | Gibco | Cat. #35050-061 | 1 mM |
| Chemical compound | Penicillin-Streptomycin | Sigma-Aldrich | Cat. # P4333 | 1% |
| Chemical compound | 4-Aminopyridine | Sigma-Aldrich | Cat. #A78403 | 100 µM |
| Chemical compound | CaCl<sub>2</sub> | Sigma-Aldrich | Cat. #C8106 | 2 mM |
| Chemical compound | MgCl<sub>2</sub> | Sigma-Aldrich | Cat. #A4425 | 1 mM |
| Chemical compound | KCl | Merk | Cat. #TA638736 | 4 mM |
| Chemical compound | ATP | Sigma-Aldrich | Cat. #A6419 | 3 mM |
| Chemical compound | GTP | Sigma-Aldrich | Cat. #G8877 | 0,1 mM |
| Chemical compound | EGTA | Sigma-Aldrich | Cat. #E4378 | 0,1 mM |
| Chemical compound | TrkB-Fc | Sigma-Aldrich | Cat. #T8694 | 1 µg/mL |
| Chemical compound | Triton X-100 | Sigma-Aldrich | Cat. #T9284 | 0,2% |
| Chemical compound | Sucrose | AppliChem | Cat. #A2211 | 4% |
| Chemical compound | Phosphate buffered saline | Sigma-Aldrich | Cat. #P4417 | 1X |
| Chemical compound | Paraformaldehyde | Sigma-Aldrich | Cat. #P6148 | 4% |
| Chemical compound | Fetal Bovine Serum, qualified, heat | Gibco | Cat. #10500064 | 5% |
| Chemical compound | Tetrodotoxin (TTX) | Tocris | Cat. #1078 | 1 μM |
| Chemical compound | D-AP5 | Tocris | Cat. #0106 | 50 μM |
| Chemical compound | CNQX | Tocris | Cat. #0190 | 10 μM |
| Chemical compound | QX314 | Tocris | Cat. # 1014 | 10 mM |
| Chemical compound | EDTA | AppliChem | Cat. #A5097 | 10 mM |
| Chemical compound | NaCl | Sigma-Aldrich | Cat. #793566 | 1 M |
| Chemical compound | Tris-HCl | Merk | Cat. # 10812846001 | 100 mM |
| Chemical compound | Tris(2-carboxyethyl)phosphine (TCEP) | Sigma-Aldrich | Cat#C4706 | 50 mM |
| Chemical compound | Protease inhibitor cocktail | Cell Signaling | Cat # #5871 | n/a |
| Chemical compound | BCA Assay | Thermo Scientific | Cat #23225 | n/a |
| Chemical compound | Hoechst 33342 | Thermo Scientific | Cat. #H1399 | 3,33 μl/mL |
| Chemical compound | TRIzol Reagent | Invitrogen | Cat. #15596018 | n/a |
| Lentiviral vectors | pcDNA6.2-GW/EmGFP-miR | Invitrogen | Cat. #K493600 | BLOCK-iT™ Pol II miR RNAi Expression Vector Kit |
| Lentiviral vectors | pCCL.sin.cPPT.PGK.GFP.WPRE | TIGET, San Raffaele Sci. Institute | gift from M. Amendola and L. Naldini | n/a |
| Nucleic compound | ODN (Positive decoy) | Sigma Genosys | Soldati et al., 2011 | n/a |
| Nucleic compound | NEG (Negative decoy) | Sigma Genosys | Soldati et al., 2011 | n/a |
| Antibodies | beta3-Tubulin (Chicken Polyclonal) | Synaptic Systems | Cat. #302306 | 1:500 |
| Antibodies | VGAT cytoplasmic domain (Mouse Monoclonal) | Synaptic Systems | Cat. #131011 | 1:500 |
| Antibodies | Gephyrin (Mouse Monoclonal) | Synaptic Systems | Cat. #147011 | 1:200 |
| Antibodies | Alexa405 goat anti-chicken | Abcam | Cat. #ab175674 | 1:500 |
| Antibodies | Alexa546 goat anti-rabbit | Thermo fisher | Cat. #A11010 | 1:500 |
| Antibodies | Alexa647 goat anti-mouse | Thermo fisher | Cat. #A32728 | 1:500 |
| Antibodies | Anti-REST (Rabbit Polyclonal) | Millipore | Cat. #07-579 | 1:500 |
| Antibodies | NPAS4 (Mouse Monoclonal) | Thermo fisher | Cat. #S408-79 | 1:500 |
| Antibodies | Synaptotagmin 4 (Rabbit Polyclonal) | Synaptic Systems | Cat. #105043 | 1:2000 |
| Antibodies | Anti-GAD67 (Mouse Monoclonal) | Sigma-Aldrich | Cat. #MAB5406 | 1:2000 |
| Antibodies | Anti-GAPDH (Rabbit Polyclonal) | Santa Cruz | Cat. #Sc-25778 | 1:2000 |
| Antibodies | Anti-mouse Peroxidase conjugated | Bio-Rad | Cat. #1706516 | 1:3000 |
| Antibodies | Anti-rabbit Peroxidase conjugated | Bio-Rad | Cat. #1706515 | 1:3000 |

### Experimental animals

GAD67-GFP knock-in mice were generated by inserting the cDNA encoding enhanced GFP into the GAD67 locus in TT2 embryonic stem cells, as described (Tamamaki et al., 2003). Heterozygous GAD67-GFP males were mated with wild-type C57BL6/J females, and GFP-positive pups were identified at birth through a Dual Fluorescent Protein Flashlight (DFP-1, NIGHTSEA, Lexington, MA USA) and confirmed by genotyping, performed by PCR with the following primers: TR-1b: GGCACAGCTCTCCCTTCTGTTTGC; TR-3:GCTCTCCTTTCGCGTTCCGACAG; TRGFP-8:CTGCTTGTCGGCCATGATATAGACG. All animals were provided by our institutional animal breeding facility in accordance with the guidelines approved by the local Animal Care Committee of the University of Genova. All experiments were carried out in accordance with the guidelines established by the European Communities Council (Directive 2010/63/EU of 4 March 2014) and were approved by the Italian Ministry of Health (authorization 73/2014-PR and 1276/2015-PR).

### Cell cultures

Primary hippocampal and cortical neurons were prepared from postnatal GAD67-GFP knock-in mice (P0–P1), as previously described (Beaudoin et al., 2012; Valente et al., 2012). Experiments used 0/2-day-old pups. Some control experiments were done using wild-type C57BL6/J mice. Briefly, hippocampi and cortex were dissociated by enzymatic digestion in 0.25% trypsin for 6 min at 37 °C and then triturated with a fire-polished Pasteur pipette. No antimitotic drugs were added to prevent glia proliferation. The following solutions were used for cell culture preparations: HANKS solution, prepared from HBSS (GIBCO 14170-088; red) supplemented with 10 mM HEPES, 30 mM D-glucose, 5 μg/ml Gentamycin, pH 7.4 with KOH; dissection solution, prepared from HANKS solution supplemented with 10% bovine serum albumin and 6 mM MgSO4. Primary hippocampal neurons were plated at density of 120 cells/mm^2^ on 3.5-cm diameter Petri dishes (Falcon® 35 mm, 353001) treated for 24 h with poly-L-lysine (0.1 mg/ml; Sigma-Aldrich) in borate buffer (0.1 M). Cells were grown in a culture medium consisting of Neurobasal A (Gibco™) supplemented with 2% B-27 (Invitrogen, Italy), 1 mM Glutamax, and 5 μg/ml Gentamycin and maintained at 37 °C in a humidified incubator with 5% CO2.

### Oligodeoxynucleotides decoy

REST activity was blocked by using oligodeoxynuclotides (ODNs) that act as surrogate binding sites for REST and sequester the native transcription factor from its genomic binding sites (Johnson et al., 2006; McClelland et al., 2011; Soldati et al., 2011). The ODN decoy (ODN) was designed corresponding to the canonical REST-binding site, RE1, while a negative decoy control (NEG) was generated using a sequence corresponding to a non-canonical RE1 that does not bind REST (Bruce et al. 2004). The decoy ODN sequences were: ODN (Positive decoy): (Top) 5’-GpPpCpPTPTTCAGCACCACGGACAGCGCCAGC-3’ (Bot) 3’-GpPpCpPTPGGCGCTGTCCGTGGTGCTGAAAGC-5’; NEG (Negative decoy) (Top) 5’-GpPpCpPTPTCCAGCACAGTGGTCAGACCC-3’ (Bot) 3’-GpPpCpPTPTCTGACCACTGTGCTGGAAGC-5’. ODNs were designed with phosphorothiolate modification on the first three nucleotides to avoid degradation (Lee et al., 2003; Osako et al., 2007). Single stranded oligodeoxynucleotides were synthesized by Sigma Genosys (St. Louis, MO). Annealing was performed in 10X buffer (100 mM Tris-HCl, pH 8.0, 10 mM EDTA, 1 M NaCl) by heating to at least 5-10 °C above their melting temperature and cooling slowly using a heat block. Cultured neurons were treated with 200 nM decoys ODNs. For the live-cell imaging of activity-dependent REST translocation from the cytoplasm to the nucleus we used the following fluorescent ODNs: ODN-Top-Cy3: 5’-GpPpCpPTP TTCAGCACCACGGACAGCGCCAGC-Cy-3; NEG-Top-Cy3: 5’-GpPpCpPTPTCCAGCACAGTGGTCAGACCC-Cy3-3’.

### RNA interference, lentivirus production and infection procedures

The REST target sequence used to design the shRNA that we employed in Figure5B was: 5’-ACATGCAAGACAGGTTCACAA-3’. This shRNA together with an alternative one were previously validated for their capability of decreasing REST mRNA levels in neuroblastoma and cultured cortical neurons (Pecoraro-Bisogni et al., 2018; Pozzi et al., 2013). Briefly, the shRNA construct was obtained by cloning the sequence into pcDNA6.2-GW/EmGFP-miR plasmid using a microRNA (miR)-based expression vector kit (BLOCK-iT Pol II miR; Invitrogen), thereby creating an expression cassette consisting of the 5′ miR flanking region, the REST target sequence, and the 3′ miR flanking region. As a negative control, the pcDNA6.2-GW/EmGFP-miR-neg plasmid (Invitrogen), containing a sequence not targeting any known vertebrate gene, was used. The shRNA was then sub-cloned into the lentiviral vector pCCL.sin.cPPT.PGK.GFP.WPRE (a kind gift from M. Amendola and L. Naldini, TIGET, San Raffaele Sci. Institute, Milan, Italy). Cultured neurons were infected at 14 div by using 10 multiplicity of infection (MOI), and neurons were used at 18-20 div.

#### Multielectrode array recordings

Neuronal network activity was recorded using a multi-well MEA system (Maestro, Axion BioSystems, Atlanta, GA). The MEA plates (M768-tMEA-48W) were composed of 48 wells, each containing a square grid of 16 PEDOT electrodes (50 μm electrode diameter; 350-μm center-to-center spacing) that create a 1.1 x 1.1 mm recording area. Spiking activity from networks grown onto MEAs was recorded and monitored using Axion BioSystems hardware (Maestro1 amplifier and Middle-man data acquisition interface) and the Axion’s Integrated Studio software in Spontaneous Neural Configuration (AxIS 2.4). After 1200x amplification, raw data were digitized at 12.5 KHz/channel and stored on a hard disk for subsequent offline analysis. The day before culture preparation, MEAs were coated by depositing a 20 μl drop of poly-L-lysine (0.1 mg/ml, Sigma-Aldrich) over each recording area and subsequently incubated overnight. After thorough washing, dissociated cortical neurons were plated at a density of 50,000 per well. Cells were incubated in Neurobasal medium supplemented with 1% Glutamax, 2% B27, 1% penicillin/streptomycin. Half-volume replacement of the culture medium was performed every 3 days. Experiments were performed in culture medium maintained at 37 °C. At div 17, MEA plates were set on the Maestro apparatus and their spontaneous activity recorded for 10 min in standard culture medium (basal condition). Cultured neurons were then treated with either ODN or NEG (200 nM) in the presence of 4AP (100 μM) or vehicle. To remove the effects of mechanical perturbations and to allow neuronal activity to reach a steady state under 4AP, MEA plates were returned to the incubator and recordings were collected 1, 24 and 48 h after the treatment. On the last day of recording, networks were acutely exposed to the GABAAR blocker bicuculline (BIC, 30 μM, Tocris Bioscience). After 5 min re-equilibration, recordings continued for 10 min (Valente et al., 2019). Spike and burst detection were both computed using the Axion BioSystems software NeuralMetricTool. To study firing and bursting properties, only the wells that contained ≥ 4 active electrodes (≥ 5 spikes/min) were retained for further analysis. Extracellular action potentials were detected by adaptive threshold crossing (7x the standard deviation of the rms-noise on each channel) on 200 Hz high-pass-filtered traces. Bursts within single channels were identified by applying an inter-spike interval (ISI) threshold algorithm (Chiappalone et al., 2006) that defines bursts as collections of a minimum number of spikes (Nmin = 5) separated by a maximum ISI (ISImax) of 100 ms. Electrodes that recorded less than 5 spikes/min were deemed inactive and were not included in the burst analysis.

#### Patch-clamp recordings

Whole-cell patch-clamp recordings were performed in the voltage-clamp configuration on hippocampal neurons plated at density of 120 cells/mm^2^. Electrophysiological experiments were performed at 17-19 div after treatment with either ODN or NEG (200 nM) in the presence of 4AP (100 μM) or vehicle for 48 h. Patch pipettes, prepared from thin borosilicate glass (Kimble, Kimax, Mexico), were pulled and fire-polished to a final resistance of 3–4 MΩ when filled with the intracellular solution. IPSCs were recorded in Tyrode extracellular solution containing (in mM): 140 NaCl, 2 CaCl2, 1 MgCl2, 4 KCl, 10 glucose, and 10 HEPES (pH 7.3 with NaOH), to which D-2-amino-5-phosphonopentanoic acid (D-AP5; 50 µM), 6-cyano-7-nitroquinoxaline-2,3-dione (CNQX; 10 µM) and N-(2,6-dimethylphenylcarbamoylmethyl)triethylammonium chloride (QX314; 10 mM) were added to block NMDA receptors, non-NMDA receptors and voltage-activated Na^+^ channels, respectively. The internal solution composition was (in mM): 140 KCl, 4 NaCl, 1 MgSO4, 0.1 EGTA, 15 glucose, 5 HEPES, 3 ATP, and 0.1 GTP (pH 7.2 with KOH). All recordings were performed at 22–24 °C. Under this condition, internal and external chloride concentrations were equimolar, shifting the chloride reversal potential from a negative value to 0 mV. This experimental configuration, that transforms IPSCs in inward currents, is typically used for increasing the amplitude of the IPSCs, evoked at negative holding potentials. IPSCs were acquired at 20 kHz sample frequency and filtered at half the acquisition rate with an 8-pole low-pass Bessel filter. Patch-clamp recordings with leak currents > 200 pA or series resistance > 15 MΩ were discarded. Series resistance was compensated 80% (2 μs response time) and the compensation was readjusted before each stimulation. The shown potentials were not corrected for the measured liquid junction potential (9 mV). Data acquisition was performed using PatchMaster and analyzed with Fit master programs (HEKA Elektronik).

For recording eIPSCs, the postsynaptic GAD67-GFP negative excitatory cell was clamped at -70mV and the soma of the presynaptic GAD67-GFP positive interneuron was stimulated through a glass electrode (1 µm tip diameter) filled with Tyrode solution in a *“loose patch configuration*”. The stimulating extracellular pipette delivered biphasic current pulses lasting 0.5 ms of variable amplitude (50–150 mA) by an isolated pulse stimulator (model 2100; A-M Systems, Carlsburg, WA). Monosynaptically connected neurons were identified by the short latency (2–4 ms) necessary to induce eIPSCs. To ensure that only the synaptic contacts of the selected presynaptic neuron were stimulated by the extracellular stimulating pipette, we recorded only those eIPSCs that were completely lost after a few µm displacements from the soma of the presynaptic neuron. Considering that the evoked currents remained stable for stimulation intensities two-fold the threshold, the stimulation intensity was set at 1.5-fold the threshold for all experiments. The current artifact produced by the presynaptic extracellular stimulation was subtracted in all the eIPSC traces. Only cells with resting membrane potentials between -57 and -64 mV were considered for analysis. eIPSCs were evoked by two consecutive stimuli separated by a time interval of 50 ms for calculating the paired-pulse ratio (PPR; I2/I1), where I1 and I2 are the amplitudes of the eIPSCs evoked by the conditioning and test stimuli, respectively. The amplitude of I2 was determined as the difference between the I2 peak and the corresponding value of I1 calculated by mono-exponential fitting of the eIPSC decay. A similar experimental protocol was used to evaluate whether BDNF played a functional role in the 4AP-induced strengthening of inhibitory synapses. To block BDNF activity, 17 div cultured neurons were incubated with TrkB-Fc (1 µg/mL, T8694-Sigma) a BDNF scavenger suppressing BDNF binding to TrkB (Sakuragi *et al*, 2013; Shelton *et al*, 1995).

The size of the RRP of synchronous release (RRPsyn) and the probability for any given synaptic vesicle in the RRP to be released (Pr) were calculated using the cumulative amplitude analysis (Schneggenburger et al., 1999). High frequency stimulation (2 s at 20 Hz) was applied to presynaptic fibers with the extracellular electrode and the RRPsyn was determined by summing up peak IPSC amplitudes during the 40 stimuli. The analysis assumes that the depression induced by the train is limited by a constant recycling of synaptic vesicles and that equilibrium is present between released and recycled vesicles. The number of data points to include in the fit of the steady-state phase was evaluated by calculating, for each cell, the best linear fit which included the maximal number of data points starting from the last one. According to this procedure, the intercept with the y-axis gives an estimation of the size of the RRPsyn and the ratio between the amplitude of the first eIPSC evoked by the stimulation train (I1) and RRPsyn yields an estimation of Pr.

Miniature IPSCs (mIPSCs) were recorded from hippocampal GAD67-GFP negative excitatory and GAD67-GFP positive inhibitory neurons incubated in the presence of tetrodotoxin (TTX, 1 μM; Tocris) to block spontaneous action potentials. mIPSC analysis was performed by using the Minianalysis program (Synaptosoft, Leonia, NJ) and the Prism software (GraphPad Software, Inc.). The amplitude and frequency of mIPSCs were calculated using a peak detector function with a threshold amplitude set at 4 pA and a threshold area at 50 ms*pA.

#### Immunocytochemistry

Immunocytochemistry of primary hippocampal neurons obtained from postnatal GAD67-GFP knock-in mice was performed after treatment with either ODN or NEG (200 nM) in the presence of 4AP (100 μM) or vehicle for 48 h. Neurons were fixed at 19 div with 4% paraformaldehyde/4% sucrose for 12 min at room temperature and then washed with phosphate buffered saline (PBS). Cells were then permeabilized with methanol (-20 °C; 10 min on ice) followed by 0.2% Triton X-100 for 10 min (Liao et al., 2010) and washed for 30 min with phosphate-buffered saline (PBS) supplemented with 5% fetal bovine serum (FBS)/0.1% bovine serum albumin (BSA). Incubation with primary antibodies in PBS/5% FBS/0.1% BSA was performed for 2 h, followed by washing with PBS and final incubation with secondary antibodies for 2 h. Neurons were immunostained with antibodies to chicken β3-Tubulin (1:500, Synaptic Systems 302306), rabbit vGAT (1:500, Synaptic Systems 131011) and mouse Gephyrin (1:200, Synaptic Systems 147011). Secondary antibodies were Alexa405 goat anti-chicken (Abcam, ab175674), Alexa546 goat anti-rabbit and Alexa647 goat anti-mouse (1:500 in all cases, Thermo fisher Cat. A11010, A32728).

Confocal images were acquired by using a 63x oil objective (N.A. 1,4) in a Leica TCS SP8 Confocal Laser Scanning Microscope (Leica Microsystem). Images were processed and analyzed using the ImageJ software (https://ima gej.nih.gov/ij/). For capturing in-focus images of objects at high magnification, multiple (25) images were taken at increasing focal distances (0.3 µm increments) and Z-stacking image processing was used to obtain a composite image with a greater depth of field than individual source images. Analysis of fluorescence intensity was performed on dendritic linear regions of interest (ROIs; 3-5 per image) of 60-160 μm in length and somatic surface ROIs (2-3 per image) of 1000-2000 μm^2^ under blind conditions. The fluorescence intensity profiles were analyzed in each ROI and putative inhibitory synaptic contacts were identified as the ROIs in which both vGAT and gephyrin average intensity profiles exceeded a threshold level (set at 3 times the background intensity). The synaptic density on dendrites was obtained by counting the total number of positive puncta divided by the length of the tubulin-positive segment (in μm). The synaptic density on the soma was obtained by counting the total number of positive puncta divided by the tubulin-positive soma area (in μm^2^). The indicated sample number (n) represents the number of coverslips collected from at least three independent neuronal preparations. From each coverslip, at least 10–15 images were acquired.

#### Live imaging of the REST translocation

Cultured hippocampal neurons (17div) from GAD67-GFP knock-in mice were treated with either vehicle or 4AP for 1 h and then incubated for 24 h with either ODN or NEG decoy tagged with Cyanine-3, (Cy3-ODN; Cy3-NEG). Five minutes before images acquisition, neurons were washed with Tyrode solution, stained with Hoechst-333342 (3.33 µl/ml) and further washed 2 times with Tyrode solution. Differential interference contrast and fluorescence images were acquired using an Olympus IX71 microscope with a 40x objective (Olympus LCPlanFI 40x) equipped with a Hamamatsu (ORCA-ER) camera and a Leica EL6000 fluorescence lamp. Somatic and nuclear Cy3-positive ROIs were drawn for each neuron to calculate the REST partition between the cytosol and the nucleus (Cy3-areanucleus/Cy3-areacyto ratio) in stimulated and control excitatory and inhibitory neurons. Images were analyzed using the ImageJ software.

#### Real-time qPCR

RNA was extracted with Triazol reagent and purified on RNeasy spin columns (Qiagen). RNA samples were quantified at 260 nm with an ND1000 Nanodrop spectrophotometer (Thermo Scientific). RNA purity was also determined by absorbance at 280 and 230 nm. All samples showed A260/280 and A260/230 ratios greater than 1.9. Reverse transcription was performed according to the manufacturer’s recommendations on 1 μg of RNA with the QuantiTect Reverse Transcription Kit (Qiagen), which includes a genomic DNA–removal step. SYBR green RT-qPCR was performed in triplicate with 10 ng of template cDNA using QuantiTect Master Mix (Qiagen) on a 7900-HT Fast Real-time System (Applied Biosystems) as previously described (Pozzi et al., 2013), using the following conditions: 5 min at 95 °C, 40 cycles of denaturation at 95 °C for 15 s, and annealing/extension at 60 °C for 30 s. Product specificity and occurrence of primer dimers were verified by melting-curve analysis. Primers were designed with Beacon Designer software (Premier Biosoft) to avoid template secondary structure and significant cross homology with other genes by BLAST search. The PCR reaction efficiency for each primer pair was calculated by the standard curve method with four serial-dilution points of cDNA. The PCR efficiency calculated for each primer set was used for subsequent analysis. All experimental samples were detected within the linear range of the assay. Gene-expression data were normalized via the multiple-internal-control-gene method (Vandesompele et al., 2002) with the GeNorm algorithm available in qBasePlus software (Biogazelle).

The control genes used were GAPDH (glyceraldehyde-3-phosphate dehydrogenase) and PPIA (peptidylprolyl isomerase), the expression of these genes was found not to be affected by the 4AP treatment. Primers sequences (5’-3’) were:

GAPDH-F: GAACATCATCCCTGCATCCA;

GAPDH-R: CCAGTGAGCTTCCCGTTCA;

PPIA-F: CACTGTCGCTTTTCGCCGCTTG;

PPIA-R: TTTCTGCTGTCTTTGGAACTTTGTCTGC;

REST-F: GAACCACCTCCCAGTATG;

REST-R: CTTCTGACAATCCTCCATAG;

BDNF cds-F: GATGCCGCAAACATGTCTATGA;

BDNF cds-R: TAATACTGTCACACACGCTCAGCTC;

BDNF (P1) F: TGGTAACCTCGCTCATTCATTAGA;

BDNF (P1) R:CCCTTCGCAATATCCGCAAAG;

BDNF (P4) F: CAAATGGAGCTTCTCGCTGAAGGC;

BDNF (P4) R:GTGGAAATTGCATGGCGGAGGTAA;

Npas4-F: AGGGTTTGCTGATGAGTTGC;

Npas4-R: CCCCTCCACTTCCATCTTC;

GABArƐ-F: TCAATGCGAAGAACACTTGG;

GABArƐ-R: AGAAGGAGACCCAGGAGAGC;

GAD67-F: CTAGGGACCCAGGGAAAG;

GAD67 -R: GTACATCTGTCATCCATCATCC;

GAD65-F: ACCAATTATGGAGCGTCACAGG;

GAD65-R: CTGAGGAGCAGCACCTTCTC;

vGAT-F: TTCAGTGCTTGGAATCTAC;

vGAT-R: TTCTCCAGAGTGAAGTCG;

Syt4-F: CCTCACTCATCGCCATCCA;

Syt4-R: GACCGCAGCTCACTCCAT.

#### Protein extraction and western blotting

Total cell lysates were obtained from 18/19 div cortical neurons treated as previously described. Neurons were lysed in lysis buffer (150 mM NaCl, 50 mM Tris, 1 mM EDTA and 1 % Triton X-100) supplemented with protease inhibitor cocktail (Cell Signaling, Danvers, MA). After 10 min of incubation, lysates were collected and clarified by centrifugation (10 min at 10,000 x g). Protein concentrations were determined by the BCA protein assay (Thermo Scientific, Waltham, MA). Sodium dodecyl sulfate polyacrylamide gel electrophoresis (SDS-PAGE) was performed according to Laemmli (1970); equivalent amounts of protein were separated on 8 % polyacrylamide gels and transferred onto nitrocellulose membranes (Whatman, St. Louis, MO). Membranes were blocked for 1 h in 5 % non-fat dry milk in Tris-buffered saline (10 mM Tris, 150 mM NaCl, pH 8.0) plus 0.1 % Triton X-100 and incubated overnight at 4 °C or for 2 h at room temperature with the following primary antibodies: anti-REST (1:500; 07-579 Millipore, MA), anti-Npas4 (1:500; S408-79 Thermo Scientific, Waltham, MA), anti-Syt4 (1:2000; 105043 Synaptic Systems, Gottingen, Germany), anti-GAD67 (1:2000; MAB5406, Sigma-Aldrich, St. Louis, MO), anti v-GAT (1:1000; 131002 Synaptic Systems, Gottingen, Germany), anti-GAPDH (1:2000; sc-25778 Santa Cruz Biotechnology, Dallas, TX). After several washes, membranes were incubated for 1 h at room temperature with peroxidase-conjugated anti-mouse (1:3000; BioRad, Hercules, CA) or anti-rabbit (1:3000; BioRad, Hercules, CA) antibodies. Bands were revealed with the ECL chemiluminescence detection system (Thermo Scientific, Waltham, MA). Immunoblots were quantified by densitometric analysis of the fluorograms (Quantity One software; Bio-Rad, Hercules, CA) obtained in the linear range of the emulsion response.

### Statistical Analysis

Data are given as means±sem for n=sample size. The normal distribution of experimental data was assessed using Kolmogorov-Smirnov normality test. To compare two normally distributed sample groups, the unpaired two-tailed Student’s *t*-test was used. To compare two sample groups that were not normally distributed, the Mann–Whitney’s *U*-test was used. To compare more than two normally distributed sample groups, we used one-or two-way ANOVA, followed by the Tukey’s or Dunnett’s multiple comparison tests. Alpha levels for all tests were 0.5% (95% confidence intervals). Statistical analysis was carried out by using the Prism software (GraphPad Software, Inc.).

## Acknowledgments

We thank dr. Yuchio Yanagawa (Gunma University, Maebashi, Japan) for kindly providing GAD67-GFP knock-in mice; drs. Anna Rocchi and Andrea Contestabile (Istituto Italiano di Tecnologia, Genova, Italy) for help and advice for RT-qPCR experiments; drs. M. Amendola and L. Naldini (TIGET, San Raffaele Sci. Institute, Milan, Italy) for kindly providing the lentiviral vectors; drs. Silvia Casagrande (Department Experimental Medicine, University of Genova) and Arta Mehilli (Istituto Italiano di Tecnologia, Genova, Italy) for help in the preparation of primary cultures; drs. Riccardo Navone and Diego Moruzzo (Istituto Italiano di Tecnologia, Genova, Italy) for help in the maintenance and genotyping of mouse strains. The study was supported by research grants from European Union EraNet Neuron 2017 SNAREopathies (to FB); Compagnia di San Paolo Torino (2017.20612 to PB and 2019.34760 to FB); IRCCS Ospedale Policlinico San Martino (Ricerca Corrente and “5x1000” to PB and FB) and Italian Ministry of University and Research (PRIN2017-A9MK4R to FB). The support of Telethon-Italy (Grant GGP19120 to FB) is also acknowledged.

## Author contributions

CP performed the electrophysiological, immunocytochemical and qPCR experiments and analyzed the data; DF performed the immunocytochemical experiments and the morphometric analysis and participated in electrophysiological experiments; AR performed the electrophysiological MEA experiments and analyzed the data; AM and FO performed the biochemical experiments and analyzed the data; PV participated in the electrophysiological experiments and in data analysis and interpretation; GL participated in data analysis and interpretation; FB supervised the research, interpreted and discussed the data, wrote the manuscript and funded research; PB supervised the research, performed analysis, interpreted and discussed the data, wrote the manuscript and funded research. All authors participated in data discussion, figure preparation and manuscript writing.

## REFERENCES

Aid T, Kazantseva A, Piirsoo M, Palm K, Timmusk T. 2007. Mouse and rat BDNF gene structure and expression revisited. J Neurosci Res 85:525–535. doi:10.1002/jnr.21139

Baldelli P, Hernandez-Guijo J-M, Carabelli V, Carbone E. 2005. Brain-derived neurotrophic factor enhances GABA release probability and nonuniform distribution of N- and P/Q-type channels on release sites of hippocampal inhibitory synapses. J Neurosci 25:3358–3368. doi:10.1523/JNEUROSCI.4227-04.2005

Baldelli P, Meldolesi J. 2015. The Transcription Repressor REST in Adult Neurons: Physiology, Pathology, and Diseases. eNeuro 2. doi:10.1523/ENEURO.0010-15.2015

Baldelli P, Novara M, Carabelli V, Hernández-Guijo JM, Carbone E. 2002. BDNF up-regulates evoked GABAergic transmission in developing hippocampus by potentiating presynaptic N- and P/Q-type Ca2+ channels signalling. Eur J Neurosci 16:2297–2310. doi:10.1046/j.1460- 9568.2002.02313.x

Ballas N, Battaglioli E, Atouf F, Andres ME, Chenoweth J, Anderson ME, Burger C, Moniwa M, Davie JR, Bowers WJ, Federoff HJ, Rose DW, Rosenfeld MG, Brehm P, Mandel G. 2001. Regulation of Neuronal Traits by a Novel Transcriptional Complex. Neuron 31:353–365. doi:10.1016/S0896-6273(01)00371-3

Ballas N, Grunseich C, Lu DD, Speh JC, Mandel G. 2005a. REST and Its Corepressors Mediate Plasticity of Neuronal Gene Chromatin throughout Neurogenesis. Cell 121:645–657. doi:10.1016/j.cell.2005.03.013

Ballas N, Grunseich C, Lu DD, Speh JC, Mandel G. 2005b. REST and Its Corepressors Mediate Plasticity of Neuronal Gene Chromatin throughout Neurogenesis. Cell 121:645–657. doi:10.1016/j.cell.2005.03.013

Beaudoin GMJ, Lee S-H, Singh D, Yuan Y, Ng Y-G, Reichardt LF, Arikkath J. 2012. Culturing pyramidal neurons from the early postnatal mouse hippocampus and cortex. Nat Protoc 7:1741–1754. doi:10.1038/nprot.2012.099

Bersten DC, Wright JA, McCarthy PJ, Whitelaw ML. 2014. Regulation of the neuronal transcription factor NPAS4 by REST and microRNAs. Biochimica et Biophysica Acta (BBA) - Gene Regulatory Mechanisms 1839:13–24. doi:10.1016/j.bbagrm.2013.11.004

Bloodgood BL, Sharma N, Browne HA, Trepman AZ, Greenberg ME. 2013. The activity-dependent transcription factor NPAS4 regulates domain-specific inhibition. Nature 503:121–125. doi:10.1038/nature12743

Brigadski T, Hartmann M, Lessmann V. 2005. Differential vesicular targeting and time course of synaptic secretion of the mammalian neurotrophins. J Neurosci 25:7601–7614. doi:10.1523/JNEUROSCI.1776-05.2005

Bruce AW, Donaldson IJ, Wood IC, Yerbury SA, Sadowski MI, Chapman M, Gottgens B, Buckley NJ. 2004. Genome-wide analysis of repressor element 1 silencing transcription factor/neuron-restrictive silencing factor (REST/NRSF) target genes. Proceedings of the National Academy of Sciences 101:10458–10463. doi:10.1073/pnas.0401827101

Canals JM, Checa N, Marco S, Akerud P, Michels A, Pérez-Navarro E, Tolosa E, Arenas E, Alberch J. 2001. Expression of brain-derived neurotrophic factor in cortical neurons is regulated by striatal target area. J Neurosci 21:117–124.

Chiappalone M, Bove M, Vato A, Tedesco M, Martinoia S. 2006. Dissociated cortical networks show spontaneously correlated activity patterns during in vitro development. Brain Res 1093:41– 53. doi:10.1016/j.brainres.2006.03.049

Chong JA, Tapia-Ramírez J, Kim S, Toledo-Aral JJ, Zheng Y, Boutros MC, Altshuller YM, Frohman MA, Kraner SD, Mandel G. 1995. REST: a mammalian silencer protein that restricts sodium channel gene expression to neurons. Cell 80:949–957. doi:10.1016/0092-8674(95)90298-8

Dean C, Liu H, Dunning FM, Chang PY, Jackson MB, Chapman ER. 2009. Synaptotagmin-IV modulates synaptic function and long-term potentiation by regulating BDNF release. Nat Neurosci 12:767–776. doi:10.1038/nn.2315

Dean C, Liu H, Staudt T, Stahlberg MA, Vingill S, Bückers J, Kamin D, Engelhardt J, Jackson MB, Hell SW, Chapman ER. 2012. Distinct subsets of Syt-IV/BDNF vesicles are sorted to axons versus dendrites and recruited to synapses by activity. J Neurosci 32:5398–5413. doi:10.1523/JNEUROSCI.4515-11.2012

Dieni S, Matsumoto T, Dekkers M, Rauskolb S, Ionescu MS, Deogracias R, Gundelfinger ED, Kojima M, Nestel S, Frotscher M, Barde Y-A. 2012. BDNF and its pro-peptide are stored in presynaptic dense core vesicles in brain neurons. J Cell Biol 196:775–788. doi:10.1083/jcb.201201038

Edwards FA, Konnerth A, Sakmann B. 1990. Quantal analysis of inhibitory synaptic transmission in the dentate gyrus of rat hippocampal slices: a patch-clamp study. J Physiol (Lond) 430:213– 249. doi:10.1113/jphysiol.1990.sp018289

Ernfors P, Wetmore C, Olson L, Persson H. 1990. Identification of cells in rat brain and peripheral tissues expressing mRNA for members of the nerve growth factor family. Neuron 5:511–526. doi:10.1016/0896-6273(90)90090-3

Fiorentino H, Kuczewski N, Diabira D, Ferrand N, Pangalos MN, Porcher C, Gaiarsa J-L. 2009. GABA(B) receptor activation triggers BDNF release and promotes the maturation of GABAergic synapses. J Neurosci 29:11650–11661. doi:10.1523/JNEUROSCI.3587-09.2009

Frere S, Slutsky I. 2018. Alzheimer’s Disease: From Firing Instability to Homeostasis Network Collapse. Neuron 97:32–58. doi:10.1016/j.neuron.2017.11.028

Garriga-Canut M, Schoenike B, Qazi R, Bergendahl K, Daley TJ, Pfender RM, Morrison JF, Ockuly J, Stafstrom C, Sutula T, Roopra A. 2006. 2-Deoxy-D-glucose reduces epilepsy progression by NRSF-CtBP-dependent metabolic regulation of chromatin structure. Nat Neurosci 9:1382–1387. doi:10.1038/nn1791

Griffith EC, Cowan CW, Greenberg ME. 2001. REST Acts through Multiple Deacetylase Complexes. Neuron 31:339–340. doi:10.1016/S0896-6273(01)00386-5

Hara D, Fukuchi M, Miyashita T, Tabuchi A, Takasaki I, Naruse Y, Mori N, Kondo T, Tsuda M. 2009. Remote control of activity-dependent BDNF gene promoter-I transcription mediated by REST/NRSF. Biochem Biophys Res Commun 384:506–511. doi:10.1016/j.bbrc.2009.05.007

Hofer M, Pagliusi SR, Hohn A, Leibrock J, Barde YA. 1990. Regional distribution of brain-derived neurotrophic factor mRNA in the adult mouse brain. EMBO J 9:2459–2464.

Hu X-L, Cheng X, Cai L, Tan G-H, Xu L, Feng X-Y, Lu T-J, Xiong H, Fei J, Xiong Z-Q. 2011. Conditional Deletion of NRSF in Forebrain Neurons Accelerates Epileptogenesis in the Kindling Model. Cereb Cortex 21:2158–2165. doi:10.1093/cercor/bhq284

Huang ZJ, Kirkwood A, Pizzorusso T, Porciatti V, Morales B, Bear MF, Maffei L, Tonegawa S. 1999. BDNF regulates the maturation of inhibition and the critical period of plasticity in mouse visual cortex. Cell 98:739–755. doi:10.1016/s0092-8674(00)81509-3

Jiao Y, Zhang Z, Zhang C, Wang X, Sakata K, Lu B, Sun Q-Q. 2011. A key mechanism underlying sensory experience-dependent maturation of neocortical GABAergic circuits in vivo. Proceedings of the National Academy of Sciences 108:12131–12136. doi:10.1073/pnas.1105296108

Johnson DS, Mortazavi A, Myers RM, Wold B. 2007. Genome-wide mapping of in vivo protein-DNA interactions. Science 316:1497–1502. doi:10.1126/science.1141319

Johnson R, Gamblin RJ, Ooi L, Bruce AW, Donaldson IJ, Westhead DR, Wood IC, Jackson RM, Buckley NJ. 2006. Identification of the REST regulon reveals extensive transposable element-mediated binding site duplication. Nucleic Acids Res 34:3862–3877. doi:10.1093/nar/gkl525

Kallunki P, Edelman GM, Jones FS. 1998. The neural restrictive silencer element can act as both a repressor and enhancer of L1 cell adhesion molecule gene expression during postnatal development. Proc Natl Acad Sci U S A 95:3233–3238. doi:10.1073/pnas.95.6.3233

Kash SF, Johnson RS, Tecott LH, Noebels JL, Mayfield RD, Hanahan D, Baekkeskov S. 1997. Epilepsy in mice deficient in the 65-kDa isoform of glutamic acid decarboxylase. Proc Natl Acad Sci USA 94:14060–14065. doi:10.1073/pnas.94.25.14060

Kuczewski N, Porcher C, Gaiarsa J-L. 2010. Activity-dependent dendritic secretion of brain-derived neurotrophic factor modulates synaptic plasticity. Eur J Neurosci 32:1239–1244. doi:10.1111/j.1460-9568.2010.07378.x

Lee IK, Ahn JD, Kim HS, Park JY, Lee KU. 2003. Advantages of the circular dumbbell decoy in gene therapy and studies of gene regulation. Curr Drug Targets 4:619–623. doi:10.2174/1389450033490821

Liao IH, Corbett BA, Gilbert DL, Bunge SA, Sharp FR. 2010. Blood gene expression correlated with tic severity in medicated and unmedicated patients with Tourette Syndrome. Pharmacogenomics 11:1733–1741. doi:10.2217/pgs.10.160

Lignani G, Baldelli P, Marra V. 2020. Homeostatic Plasticity in Epilepsy. Front Cell Neurosci 14:197. doi:10.3389/fncel.2020.00197

Lin TW, Harward SC, Huang YZ, McNamara JO. 2020. Targeting BDNF/TrkB pathways for preventing or suppressing epilepsy. Neuropharmacology 167:107734. doi:10.1016/j.neuropharm.2019.107734

Lin Y, Bloodgood BL, Hauser JL, Lapan AD, Koon AC, Kim T-K, Hu LS, Malik AN, Greenberg ME. 2008. Activity-dependent regulation of inhibitory synapse development by Npas4. Nature 455:1198–1204. doi:10.1038/nature07319

Lu T, Aron L, Zullo J, Pan Y, Kim H, Chen Y, Yang T-H, Kim H-M, Drake D, Liu XS, Bennett DA, Colaiácovo MP, Yankner BA. 2014. REST and stress resistance in ageing and Alzheimer’s disease. Nature 507:448–454. doi:10.1038/nature13163

Marty S, Wehrlé R, Sotelo C. 2000. Neuronal activity and brain-derived neurotrophic factor regulate the density of inhibitory synapses in organotypic slice cultures of postnatal hippocampus. J Neurosci 20:8087–8095.

Matsumoto T, Rauskolb S, Polack M, Klose J, Kolbeck R, Korte M, Barde Y-A. 2008. Biosynthesis and processing of endogenous BDNF: CNS neurons store and secrete BDNF, not pro-BDNF. Nat Neurosci 11:131–133. doi:10.1038/nn2038

McClelland S, Brennan GP, Dubé C, Rajpara S, Iyer S, Richichi C, Bernard C, Baram TZ. 2014. The transcription factor NRSF contributes to epileptogenesis by selective repression of a subset of target genes. Elife 3:e01267. doi:10.7554/eLife.01267

McClelland S, Flynn C, Dubé C, Richichi C, Zha Q, Ghestem A, Esclapez M, Bernard C, Baram TZ. 2011. Neuron-restrictive silencer factor-mediated hyperpolarization-activated cyclic nucleotide gated channelopathy in experimental temporal lobe epilepsy. Ann Neurol 70:454– 464. doi:10.1002/ana.22479

O’Brien RJ, Kamboj S, Ehlers MD, Rosen KR, Fischbach GD, Huganir RL. 1998. Activity-dependent modulation of synaptic AMPA receptor accumulation. Neuron 21:1067–1078. doi:10.1016/s0896-6273(00)80624-8

Ohba S, Ikeda T, Ikegaya Y, Nishiyama N, Matsuki N, Yamada MK. 2005. BDNF locally potentiates GABAergic presynaptic machineries: target-selective circuit inhibition. Cereb Cortex 15:291–298. doi:10.1093/cercor/bhh130

Ooi L, Wood IC. 2007. Chromatin crosstalk in development and disease: lessons from REST. Nature Reviews Genetics 8:544–554. doi:10.1038/nrg2100

Osako MK, Tomita N, Nakagami H, Kunugiza Y, Yoshino M, Yuyama K, Tomita T, Yoshikawa H, Ogihara T, Morishita R. 2007. Increase in nuclease resistance and incorporation of NF-kappaB decoy oligodeoxynucleotides by modification of the 3’-terminus. J Gene Med 9:812– 819. doi:10.1002/jgm.1077

Otto SJ, McCorkle SR, Hover J, Conaco C, Han J-J, Impey S, Yochum GS, Dunn JJ, Goodman RH, Mandel G. 2007. A New Binding Motif for the Transcriptional Repressor REST Uncovers Large Gene Networks Devoted to Neuronal Functions. Journal of Neuroscience 27:6729– 6739. doi:10.1523/JNEUROSCI.0091-07.2007

Palm K, Belluardo N, Metsis M, Timmusk T. 1998. Neuronal expression of zinc finger transcription factor REST/NRSF/XBR gene. J Neurosci 18:1280–1296.

Paonessa F, Criscuolo S, Sacchetti S, Amoroso D, Scarongella H, Pecoraro Bisogni F, Carminati E, Pruzzo G, Maragliano L, Cesca F, Benfenati F. 2016. Regulation of neural gene transcription by optogenetic inhibition of the RE1-silencing transcription factor. Proceedings of the National Academy of Sciences 113:E91–E100. doi:10.1073/pnas.1507355112

Paonessa F, Latifi S, Scarongella H, Cesca F, Benfenati F. 2013. Specificity Protein 1 (Sp1)-dependent Activation of the Synapsin I Gene (*SYN1*) Is Modulated by RE1-silencing Transcription Factor (REST) and 5′-Cytosine-Phosphoguanine (CpG) Methylation. J Biol Chem 288:3227–3239. doi:10.1074/jbc.M112.399782

Pattabiraman PP, Tropea D, Chiaruttini C, Tongiorgi E, Cattaneo A, Domenici L. 2005. Neuronal activity regulates the developmental expression and subcellular localization of cortical BDNF mRNA isoforms in vivo. Mol Cell Neurosci 28:556–570. doi:10.1016/j.mcn.2004.11.010

Pecoraro-Bisogni F, Lignani G, Contestabile A, Castroflorio E, Pozzi D, Rocchi A, Prestigio C, Orlando M, Valente P, Massacesi M, Benfenati F, Baldelli P. 2018. REST-Dependent Presynaptic Homeostasis Induced by Chronic Neuronal Hyperactivity. Mol Neurobiol 55:4959–4972. doi:10.1007/s12035-017-0698-9

Perera A, Eisen D, Wagner M, Laube SK, Künzel AF, Koch S, Steinbacher J, Schulze E, Splith V, Mittermeier N, Müller M, Biel M, Carell T, Michalakis S. 2015. TET3 is recruited by REST for context-specific hydroxymethylation and induction of gene expression. Cell Rep 11:283–294. doi:10.1016/j.celrep.2015.03.020

Pozzi D, Lignani G, Ferrea E, Contestabile A, Paonessa F, D’Alessandro R, Lippiello P, Boido D, Fassio A, Meldolesi J, Valtorta F, Benfenati F, Baldelli P. 2013. REST/NRSF-mediated intrinsic homeostasis protects neuronal networks from hyperexcitability. EMBO J 32:2994– 3007. doi:10.1038/emboj.2013.231

Pruunsild P, Sepp M, Orav E, Koppel I, Timmusk T. 2011. Identification of cis-elements and transcription factors regulating neuronal activity-dependent transcription of human BDNF gene. J Neurosci 31:3295–3308. doi:10.1523/JNEUROSCI.4540-10.2011

Sakuragi S, Tominaga-Yoshino K, Ogura A. 2013. Involvement of TrkB- and p75(NTR)-signaling pathways in two contrasting forms of long-lasting synaptic plasticity. Sci Rep 3:3185. doi:10.1038/srep03185

Schneggenburger R, Meyer AC, Neher E. 1999. Released Fraction and Total Size of a Pool of Immediately Available Transmitter Quanta at a Calyx Synapse. Neuron 23:399–409. doi:10.1016/S0896-6273(00)80789-8

Schoenherr CJ, Anderson DJ. 1995. Silencing is golden: negative regulation in the control of neuronal gene transcription. Current Opinion in Neurobiology 5:566–571. doi:10.1016/0959- 4388(95)80060-3

Seil FJ, Drake-Baumann R. 2000. TrkB receptor ligands promote activity-dependent inhibitory synaptogenesis. J Neurosci 20:5367–5373.

Shao L-R, Dudek FE. 2005. Changes in mIPSCs and sIPSCs after kainate treatment: evidence for loss of inhibitory input to dentate granule cells and possible compensatory responses. J Neurophysiol 94:952–960. doi:10.1152/jn.01342.2004

Shelton DL, Sutherland J, Gripp J, Camerato T, Armanini MP, Phillips HS, Carroll K, Spencer SD, Levinson AD. 1995. Human trks: molecular cloning, tissue distribution, and expression of extracellular domain immunoadhesins. J Neurosci 15:477–491.

Shimojo M. 2008. Huntingtin regulates RE1-silencing transcription factor/neuron-restrictive silencer factor (REST/NRSF) nuclear trafficking indirectly through a complex with REST/NRSF-interacting LIM domain protein (RILP) and dynactin p150 Glued. J Biol Chem 283:34880– 34886. doi:10.1074/jbc.M804183200

Shimojo M, Hersh LB. 2003. REST/NRSF-interacting LIM domain protein, a putative nuclear translocation receptor. Mol Cell Biol 23:9025–9031. doi:10.1128/mcb.23.24.9025-9031.2003

Shimojo M, Lee JH, Hersh LB. 2001. Role of zinc finger domains of the transcription factor neuron-restrictive silencer factor/repressor element-1 silencing transcription factor in DNA bindingand nuclear localization. J Biol Chem 276:13121–13126. doi:10.1074/jbc.M011193200

Soldati C, Bithell A, Conforti P, Cattaneo E, Buckley NJ. 2011. Rescue of gene expression by modified REST decoy oligonucleotides in a cellular model of Huntington’s disease. J Neurochem 116:415–425. doi:10.1111/j.1471-4159.2010.07122.x

Spencer EM, Chandler KE, Haddley K, Howard MR, Hughes D, Belyaev ND, Coulson JM, Stewart JP, Buckley NJ, Kipar A, Walker MC, Quinn JP. 2006. Regulation and role of REST and REST4 variants in modulation of gene expression in in vivo and in vitro in epilepsy models. Neurobiol Dis 24:41–52. doi:10.1016/j.nbd.2006.04.020

Spiegel I, Mardinly AR, Gabel HW, Bazinet JE, Couch CH, Tzeng CP, Harmin DA, Greenberg ME. 2014. Npas4 regulates excitatory-inhibitory balance within neural circuits through cell-type-specific gene programs. Cell 157:1216–1229. doi:10.1016/j.cell.2014.03.058

Styr B, Slutsky I. 2018. Imbalance between firing homeostasis and synaptic plasticity drives early-phase Alzheimer’s disease. Nature Neuroscience 21:463–473. doi:10.1038/s41593-018- 0080-x

Sun X, Lin Y. 2017. Npas4: Linking Neuronal Activity to Memory 21.

Tamamaki N, Yanagawa Y, Tomioka R, Miyazaki J-I, Obata K, Kaneko T. 2003. Green fluorescent protein expression and colocalization with calretinin, parvalbumin, and somatostatin in the GAD67-GFP knock-in mouse. J Comp Neurol 467:60–79. doi:10.1002/cne.10905

Turrigiano GG. 2008. The self-tuning neuron: synaptic scaling of excitatory synapses. Cell 135:422– 435. doi:10.1016/j.cell.2008.10.008

Turrigiano GG, Leslie KR, Desai NS, Rutherford LC, Nelson SB. 1998. Activity-dependent scaling of quantal amplitude in neocortical neurons. Nature 391:892–896. doi:10.1038/36103

Valente P, Casagrande S, Nieus T, Verstegen AMJ, Valtorta F, Benfenati F, Baldelli P. 2012. Site-specific synapsin I phosphorylation participates in the expression of post-tetanic potentiation and its enhancement by BDNF. J Neurosci 32:5868–5879. doi:10.1523/JNEUROSCI.5275- 11.2012

Valente P, Orlando M, Raimondi A, Benfenati F, Baldelli P. 2016. Fine Tuning of Synaptic Plasticity and Filtering by GABA Released from Hippocampal Autaptic Granule Cells. Cereb Cortex 26:1149–1167. doi:10.1093/cercor/bhu301

Valente P, Romei A, Fadda M, Sterlini B, Lonardoni D, Forte N, Fruscione F, Castroflorio E, Michetti C, Giansante G, Valtorta F, Tsai J-W, Zara F, Nieus T, Corradi A, Fassio A, Baldelli P, Benfenati F. 2019. Constitutive Inactivation of the PRRT2 Gene Alters Short-Term Synaptic Plasticity and Promotes Network Hyperexcitability in Hippocampal Neurons. Cerebral Cortex 29:2010–2033. doi:10.1093/cercor/bhy079

Vandesompele J, De Preter K, Pattyn F, Poppe B, Van Roy N, De Paepe A, Speleman F. 2002. Accurate normalization of real-time quantitative RT-PCR data by geometric averaging of multiple internal control genes. Genome Biol 3:RESEARCH0034. doi:10.1186/gb-2002-3-7-research0034

Yamada MK, Nakanishi K, Ohba S, Nakamura T, Ikegaya Y, Nishiyama N, Matsuki N. 2002. Brain-derived neurotrophic factor promotes the maturation of GABAergic mechanisms in cultured hippocampal neurons. J Neurosci 22:7580–7585.

Zullo JM, Drake D, Aron L, O’Hern P, Dhamne SC, Davidsohn N, Mao C-A, Klein WH, Rotenberg A, Bennett DA, Church GM, Colaiácovo MP, Yankner BA. 2019. Regulation of lifespan by neural excitation and REST. Nature 574:359–364. doi:10.1038/s41586-019-1647-8

